# Topoisomerase 1 dependent R-loop deficiency drives accelerated replication and genomic instability

**DOI:** 10.1101/2020.07.21.214700

**Authors:** Dan Sarni, Alon Shtrikman, Yifat S. Oren, Batsheva Kerem

**Author notes:** Corresponding Author: Prof. Batsheva Kerem, Department of Genetics, The Life Science Institute The Hebrew University, Jerusalem, Israel 91904.

## Abstract

DNA replication is a complex process that is tightly regulated to ensure faithful genome duplication, and its perturbation leads to DNA damage and genomic instability. Replication stress is commonly associated with slow and stalled replication forks. Recently, accelerated replication has emerged as a non-canonical form of replication stress. However, the molecular basis underlying fork acceleration is largely unknown. Here we show that increased topoisomerase 1 (TOP1) expression induces aberrant replication fork acceleration and DNA damage by decreasing RNA-DNA hybrids (R-loops). Degradation of R-loops by overexpression of RNaseH1 also accelerates replication and generates DNA damage. Furthermore, upregulation of TOP1 by activation of the mutated HRAS oncogene leads to fork acceleration and DNA damage in pre-senescent cells. In these cells, restoration of TOP1 expression level or mild replication inhibition rescues the perturbed replication and reduces DNA damage. These findings highlight the importance of TOP1 equilibrium in the regulation of R-loop homeostasis to ensure faithful DNA replication and genome integrity.

## Introduction

DNA replication is a complex process that is tightly regulated to ensure faithful duplication of the genome. Various factors are involved in regulating the different stages of replication, including origin licensing and firing, replication elongation rate and termination (Conti et al., 2007; Fragkos et al., 2015). Under conditions that slow or even stall replication fork progression (defined as replication stress) dormant origins are activated to allow completion of DNA synthesis to maintain genome integrity (Courbet et al., 2008; Ge et al., 2007). However, insufficient compensation of the perturbed DNA replication may lead to genome instability (Gaillard et al., 2015; Zeman and Cimprich, 2014). Several factors are thought to lead to replication stress, among them are nucleotide deficiency, accumulation of RNA-DNA hybrids and DNA lesions (Gaillard et al., 2015; Zeman and Cimprich, 2014), all of which result in perturbed replication dynamics and increase genomic instability.

Genomic instability is an important hallmark of cancer and a driver of tumorigenesis (Hanahan and Weinberg, 2011; Negrini et al., 2010). In recent years, aberrant activation of several oncogenes and tumor suppressor genes was found to induce replication stress leading to accumulation of DNA damage and an increased tumorigenicity potential (Bartkova et al., 2006; Bester et al., 2011; Dominguez-Sola et al., 2007; Galanos et al., 2016; Di Micco et al., 2006). This stress was characterized by slow replication rates, fork stalling, activation of dormant origins and even re-replication. Recently, however, several studies have found accelerated replication rates following alterations in expression of various genes. These include overexpression of the oncogene Spi-1 (Rimmele et al., 2010) and the cancer associated gene ISG15 (Raso et al., 2020), down-regulation of mRNA biogenesis genes involved in mRNA processing and export (Bhatia et al., 2014; Domínguez-Sánchez et al., 2011), depletion of origin firing factors (Sedlackova et al., 2020; Zhong et al., 2013), and even inhibition of poly(ADP-ribose) polymerase (PARP) (Maya-Mendoza et al., 2018; Sugimura et al., 2008). In most of these studies accelerated replication was accompanied by DNA damage. However, whether the accelerated replication rate induces DNA damage per se and the molecular mechanism/s underlying fork acceleration are largely unknown.

Here we demonstrate the importance of TOP1 expression level in the regulation of R-loop homeostasis to ensure faithful DNA replication and genome integrity. Our study shows that increased topoisomerase 1 (TOP1) expression causes aberrant replication fork acceleration and DNA damage by decreasing R-loop levels. Furthermore, degradation of R-loops by overexpression of RNaseH1 accelerates the replication rate and generates DNA damage. Activation of the mutated HRAS (RAS) oncogene in pre-senescent cells increases TOP1 levels leading to replication fork acceleration and thereby inducing DNA damage and genomic instability. Restoration of TOP1 expression level or mild replication inhibition rescues the perturbed replication and reduces DNA damage. Altogether, these results highlight the important role of TOP1 in maintaining genome stability by controlling R-loop homeostasis, enabling tight regulation of DNA replication fork progression. Furthermore, the results reveal a novel mechanism of oncogene-induced DNA damage induced by aberrant replication fork acceleration.

## Results

### RAS expression induces replication acceleration in pre-senescent cells

RAS proteins are members of a GTP-binding protein family (Pylayeva-Gupta et al., 2011), which regulates numerous cellular processes including cell cycle progression (Downward, 2003). Mutated RAS expression induces genomic instability (Denko et al., 1994; Saavedra et al., 2000; Yang et al., 2013) leading to senescence, a cell cycle arrest state serving as an antitumor barrier. However, cells escaping this proliferation inhibition drive tumorigenesis (Halazonetis et al., 2008). Therefore, we first investigated the effect of RAS on replication-induced genomic instability in pre-senescent cells. For this, immortalized human foreskin fibroblasts were retrovirally infected with an inducible ER:HRAS-G12V vector (referred to henceforth as RAS). RAS selective expression following 4-hydroxytamoxifen (4-OHT) supplementation was verified by Western blot (Figure 1A). Following RAS activation cells entered a hyperproliferative phase, as indicated by increased population doubling and IdU incorporation already at day 2 (Figures S1A-S1C). This was followed by a decline in the proliferative potential until proliferation ceased by day 10, when cells entered senescence as indicated by reduced population doublings, reduced IdU incorporation and increased senescence associated β-gal activity (Figures S1A-S1E). Hence, the effect of RAS activation on replication dynamics and genome stability was investigated in pre-senescence RAS expressing cells up to 5 days following RAS induction.

**Figure 1.**
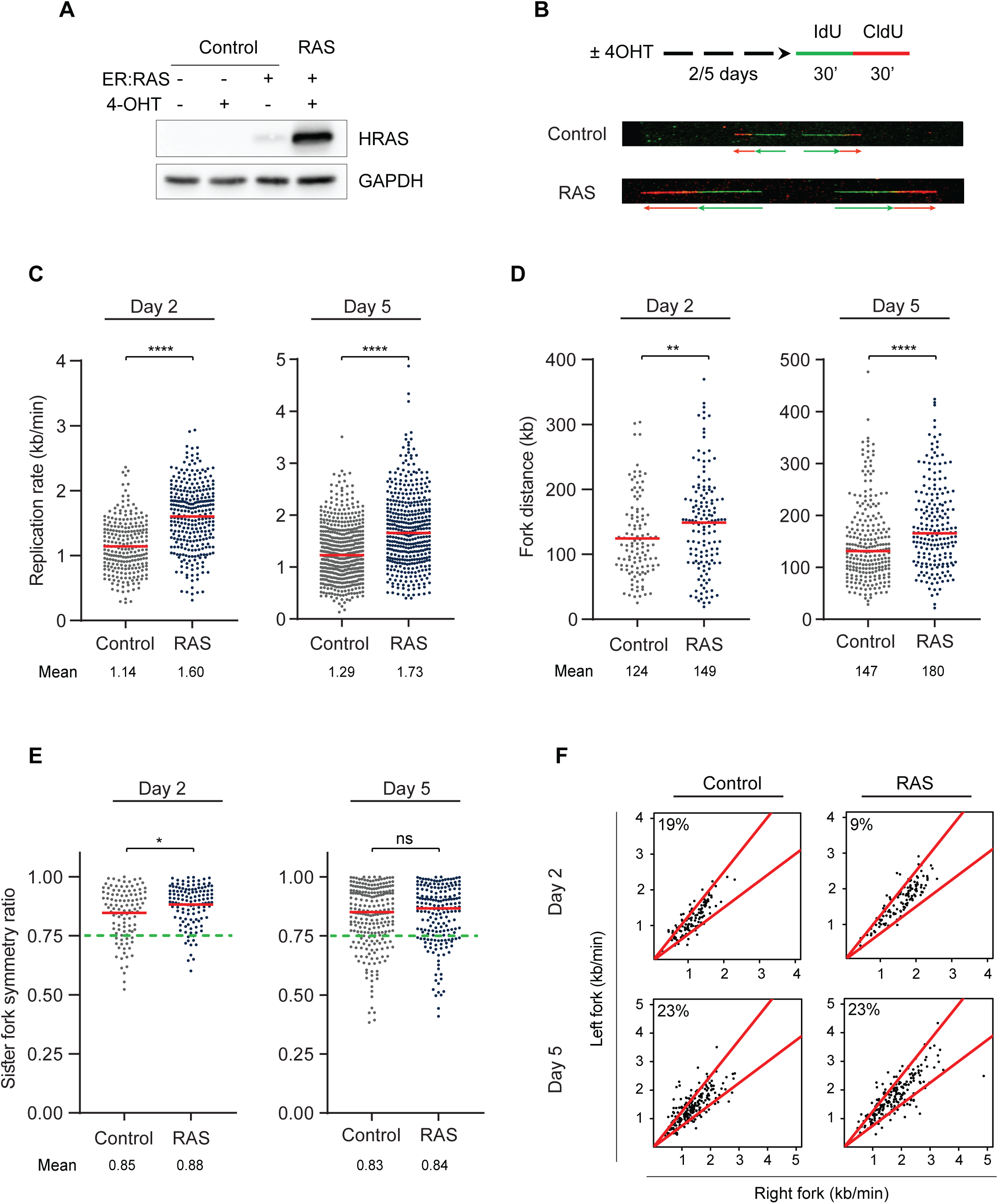
RAS expression leads to increased replication rate and fork distance. (**A**) Protein levels of HRAS and GAPDH in FSE-hTert cells with (+) or without (-) ER:RAS infection and 4- OHT treatment as indicated. (**B**) Top: a scheme of the protocol. Cells infected with ER:RAS with (RAS) or without (Control) 4-OHT treatment were pulse-labeled with two thymidine analogs (IdU then CldU) for 30 min each. DNA combing was performed 2 or 5 days after RAS induction (4-OHT treatment). Bottom: representative images of labeled DNA fibers. (**C**) Dot plots of individual replication fork rates (kb/min) in control and RAS expressing cells for 2 or 5 days; at least 240 fibers per condition were analyzed. (**D**) Dot plots of individual fork distances (kb) in control and RAS expressing cells for 2 or 5 days; at least 110 fibers per condition were analyzed. (**E**) Dot plots of sister fork symmetry ratios in control and RAS expressing cells for 2 or 5 days; at least 100 fibers per condition were analyzed. Fork symmetry is expressed as the ratio of the shorter to the longer distance covered during the IdU pulse, for each pair of sister replication forks. Dashed green line indicates asymmetry ratio threshold. (**C-E**) Red lines indicate medians, Means are indicated. (**F**) Scatter plots of the replication rates (kb/min) of right- and left-moving sister forks during IdU pulse. The center areas delimited with red lines contain sister forks with less than 25% difference. The percentages of asymmetric forks are indicated at the top left corner of the plots, control and RAS expressing cells for 2 or 5 days. (**C-F**) Data for RAS day 2 are the summary of 2 independent experiments; data for RAS day 5 are the summary of 4 independent experiments. Mann-Whitney rank-sum test, ns – non-significant; * *P* < 0.05, ** *P* < 0.01, **** *P* < 0.0001.

We then analyzed the effect of RAS activation on replication dynamics using the DNA combing approach, which enables replication analysis on single DNA molecules. Newly synthesized DNA extracted from RAS-infected cells with (RAS) or without (control) 4-OHT treatment was labeled with IdU and CldU and detected by fluorescent antibodies (green and red, respectively) (Figure 1B). The analyses were performed at two time points prior to senescence onset; at the hyperproliferative phase (day 2 post RAS activation) and at day 5 post RAS activation. First, we analyzed the effect of RAS activation on the DNA replication fork rate. The results showed a remarkable increase in the mean replication fork rate in both day 2 and day 5 following RAS activation (Figure 1C). Similar results were obtained in an additional cell line of fetal human lung fibroblasts, WI-38 (Figure S2A and S2B).

Previous studies have shown that slow replication fork progression is correlated with an increased number of activated origins (Courbet et al., 2008; Ge et al., 2007). Therefore, we investigated origin activation in cells expressing RAS in which the replication rate is accelerated. Analysis of the mean replication fork distance showed a significant increase in the mean fork distance in both day 2 and day 5 following RAS activation (Figure 1D). Similar results were obtained in WI-38 cells (Figure S2C). These results indicate that in pre-senescent RAS expressing cells there was a significant increase in the rate of replication fork progression along with a decrease in local origin activation.

Under replication stress which leads to fork stalling, asymmetrical progression of sister forks emanating from the same origin has been observed (Conti et al., 2007). In order to further characterize the accelerated replication rate following RAS expression, we analyzed the symmetry of fork progression by comparing the progression of left and right outgoing sister replication forks. As previously suggested (Bester et al., 2011; Conti et al., 2007), the threshold for asymmetric progression of two forks was considered when the minimum to maximum sister forks ratio was < 0.75. The analysis showed that most forks are symmetric in both day 2 and 5 following RAS expression as well as in control cells (Figures 1E and 1F). Similar results were obtained in WI-38 cells (Figures S2D and S2E), indicating no increase in fork stalling. Altogether, these results indicate a non-classical form of aberrant replication dynamics in pre-senescent RAS expressing cells, in which the replication fork speed is accelerated, and fewer origins are activated. This aberrant accelerated replication was observed both when RAS-cells are hyperproliferating at day 2, and at day 5 while still proliferating but shortly before they senesce.

### RAS expression leads to accelerated replication-induced DNA damage

Slow replication rate, induced by oncogenes, including mutated RAS overexpression, generates DNA damage and activates damage response pathways (Abulaiti et al., 2006; Aird et al., 2013; Kotsantis et al., 2016; Macheret and Halazonetis, 2015). In order to determine whether RAS-induced fork acceleration rate causes DNA damage we examined cellular DNA damage response markers known also to be induced under replication stress (Ewald et al., 2007). We first analyzed DNA damage formation as indicated by the co-localization of phosphorylated H2AX (γH2AX) and 53BP1 foci, after RAS activation. The analysis showed a significant increase in DNA damage in RAS-cells already at day 2 post RAS activation (Figures 2A and 2B), when aberrant acceleration of the DNA replication rate is already found (Figure 1C). Interestingly, the levels of DNA damage markers increased in RAS-cells over time (Figures 2A and 2B), implying that the accumulation of unrepaired damage may lead to cell cycle arrest, as previously suggested (Bartkova et al., 2006; Halazonetis et al., 2008; Di Micco et al., 2006). Next, we investigated whether the observed damage is associated with replication stress. Replication stress is known to induce DNA lesions which manifest as nuclear bodies of 53BP1 in the G1-phase of the next cell cycle (Lukas et al., 2011). Analysis of 53BP1 foci in G1-phase cells (cyclin A negative) showed a significant increase in foci formation in pre-senescent RAS expressing cells compared with control cells (Figures 2C and 2D). Similarly to 53BP1 foci analysis, we found in RAS expressing cells a significant increase in γH2AX, a DNA damage marker also induced upon replication stress (Figures 2E and 2F) (Ewald et al., 2007). Next, we analyzed whether RAS activation leads to chromosomal fragility, since under replication stress conditions genomic instability is found preferentially at genomic regions known as fragile sites (Glover et al., 1984; Miron et al., 2015). Metaphase spread analysis showed a significant increase in chromosomal fragility in RAS expressing cells compared with control cells (Figures 2G and 2H), implying that the replication perturbation induced by RAS generates replication stress. We then tested CHK1 activation, a hallmark of replication stress response (Zeman and Cimprich, 2014), by analyzing its phosphorylation. As can be seen in Figure 2I, a significant increase in CHK1 phosphorylation was found in RAS expressing cells compared with control cells. Thus overall, these results suggest that the RAS-induced DNA damage is associated with the aberrant acceleration of DNA replication.

**Figure 2.**
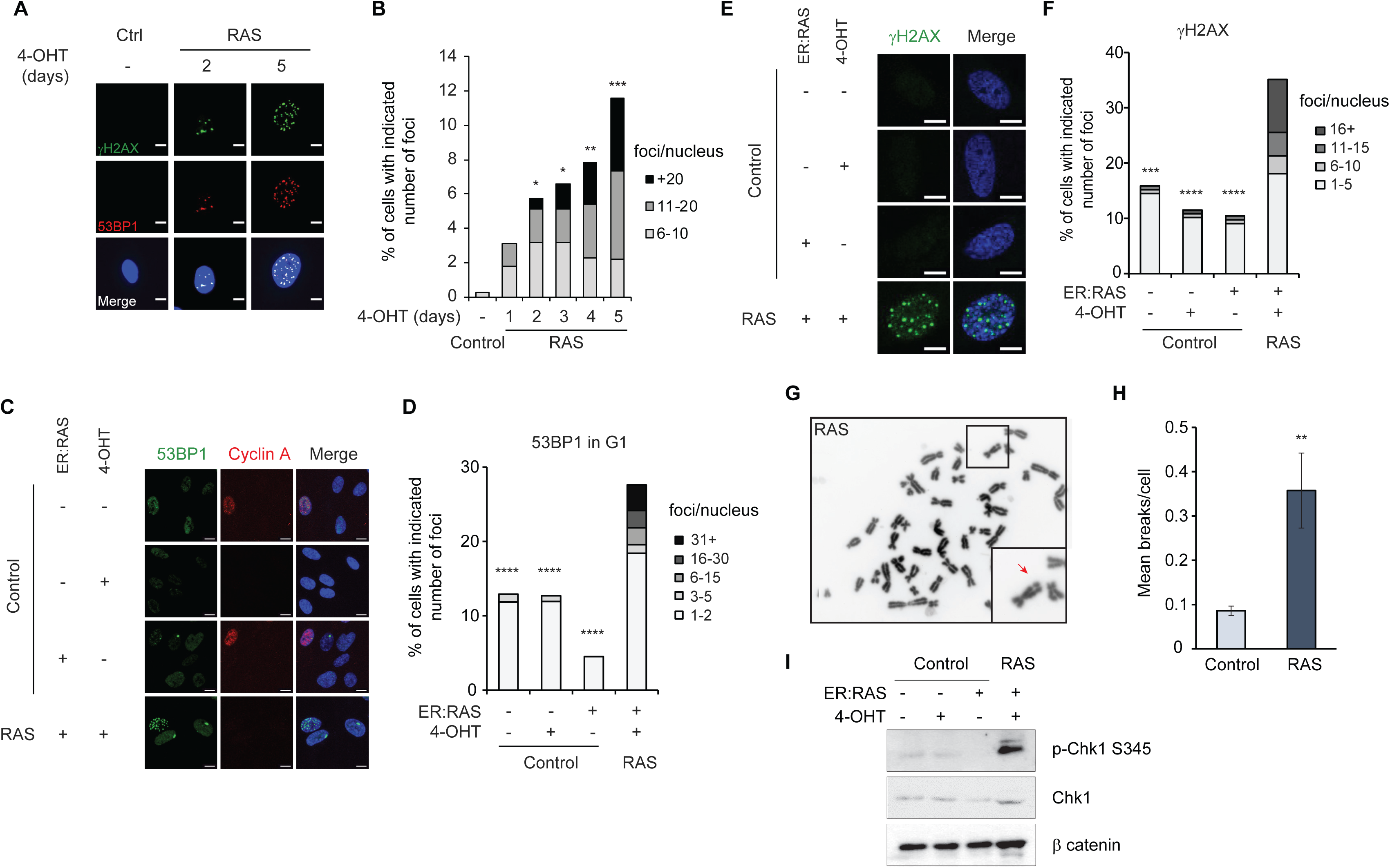
RAS expression leads to accelerated replication-induced DNA damage in pre-senescent cells. (**A,B**) Co-localization of γH2AX (red) and 53BP1 (green) foci in ER:RAS FSE-hTert cells with (+) or without (-) 4-OHT treatment for the indicated time points (days). Representative images (**A**), quantification of the percent of cells with indicated number of co- localized γH2AX and 53BP1 foci per nucleus (**B**); number of analyzed cells from left to right: n = 399, 384, 467, 762, 704, 501, respectively. Data are the summary of three independent experiments. *P* values calculated compared to ER:RAS FSE-hTert cells without 4-OHT (-) treatment (Control) by one-way ANOVA. (**C,D**) 53BP1 foci in G1-phase (cyclin A negative cells) in FSE-hTert cells with (+) or without (-) ER:RAS infection and 4-OHT treatment as indicated. Representative images of 53BP1 foci (green) in cyclin A (red) negative cells and DAPI staining (blue) (**C**), quantification of the percent of cells with indicated number of foci per nucleus (**D**); number of analyzed nuclei from left to right: n = 93, 126, 133, 87, respectively. *P* values calculated compared to ER:RAS FSE-hTert cells treated with 4-OHT (RAS). Data are representative of three independent experiments with similar results. (**E,F**) γH2AX foci in FSE- hTert cells with (+) or without (-) ER:RAS infection and 4-OHT treatment as indicated. Representative images of γH2AX foci (green) and DAPI staining (blue) (**E**), quantification of the percent of cells with the indicated number of γH2AX foci (**F**); number of analyzed nuclei from left to right: n = 145, 148, 144, 94, respectively. *P* values calculated compared to ER:RAS FSE- hTert cells treated with 4-OHT (RAS). Data are representative of two independent experiments with similar results. (**G**) A representative image of a metaphase spread in FSE-hTert RAS cells, the box at the bottom right corner shows magnification of the chromosomes in the selected smaller box; red arrow indicates a break. (**H**) Quantification of chromosomal aberrations detected in metaphase spreads of ER:RAS FSE-hTert cells treated with 4-OHT for 4 days (RAS, n = 120) or without 4-OHT treatment (Control, n = 155). The results are the average number of chromosomal aberrations from two independent experiments, means ± s.e.m are shown. (**I**) Protein levels of phosphorylated Chk1 ser345, Chk1 and β-catenin in FSE-hTert cells with (+) or without (-) ER:RAS infection and 4-OHT treatment as indicated. Mann Whitney rank-sum test, \**P* < 0.05, ** *P* < 0.01, *** *P* < 0.001; **** *P* < 0.0001. Scale bars, 10 μm.

Previously, we showed that oncogene-induced replication stress resulted from nucleotide insufficiency (Bester et al., 2011). Viral and cellular oncogenes were shown to enforce cell proliferation without sufficient nucleotide biosynthesis to support normal replication (Bester et al., 2011). Therefore, we next investigated whether the replication-induced DNA damage following RAS expression could also result from a nucleotide deficiency. We analyzed 53BP1 foci in the G1-phase of RAS expressing cells grown in a regular medium or supplemented with exogenous nucleosides. The analysis showed no significant difference in replication-induced DNA damage foci (Figure S3), indicating that DNA damage induced by fork acceleration was not the result of nucleotide insufficiency.

### Reducing the accelerated replication fork progression rescues the DNA damage

We next investigated whether the replication fork acceleration could be the cause for DNA damage formation in RAS expressing pre-senescent cells. For this, we slowed down the replication fork progression in RAS-cells by treating RAS expressing cells with hydroxyurea (HU), a known inhibitor of replication fork progression and analyzed its effect on DNA damage formation. HU inhibits the ribonucleotide reductase (RNR), thus reducing the deoxyribonucleotide pool, resulting in reduced replication fork progression in a dose-dependent manner (Ge and Blow, 2010; Skoog and Nordenskjöld, 1971; Técher et al., 2016). Whereas high doses of HU (≥1mM) lead to fork arrest, low doses (≤ 0.1mM) decelerate replication fork progression (Koundrioukoff et al., 2013; Técher et al., 2016). Therefore, we used various relatively low HU concentrations and analyzed their effect on the replication dynamics. Cells expressing RAS for 5 days were treated with 0.001-0.1mM HU for 48 hours prior to the analysis, which allowed the cells to go through at least one cell cycle under inhibitory conditions (Figure 3A). Flow cytometry analysis showed no significant change in the cell cycle progression, indicating that HU treatment did not arrest cell proliferation (Figures S4A and S4B). DNA combing analysis revealed that mild replication inhibition of RAS-cells with 0.01mM HU resulted in a dramatic replication deceleration compared with non-treated RAS-cells (Figure 3B). The mean rate in these HU-treated RAS-cells showed no significant difference compared with control cells, indicating restoration of a normal replication fork rate (Figure 3B). This HU concentration also led to a reduction in the fork distance in RAS expressing cells to the normal distance observed in the control cells (Figure 3C). Finally, 0.01mM HU treatment did not induce sister fork asymmetry in RAS expressing cells, indicating that this low HU concentration did not induce fork stalling (Figure 3D). Thus overall, these results indicate that mild replication inhibition in RAS-cells rescued the perturbed DNA replication, resulting in the restoration of normal replication dynamics.

**Figure 3.**
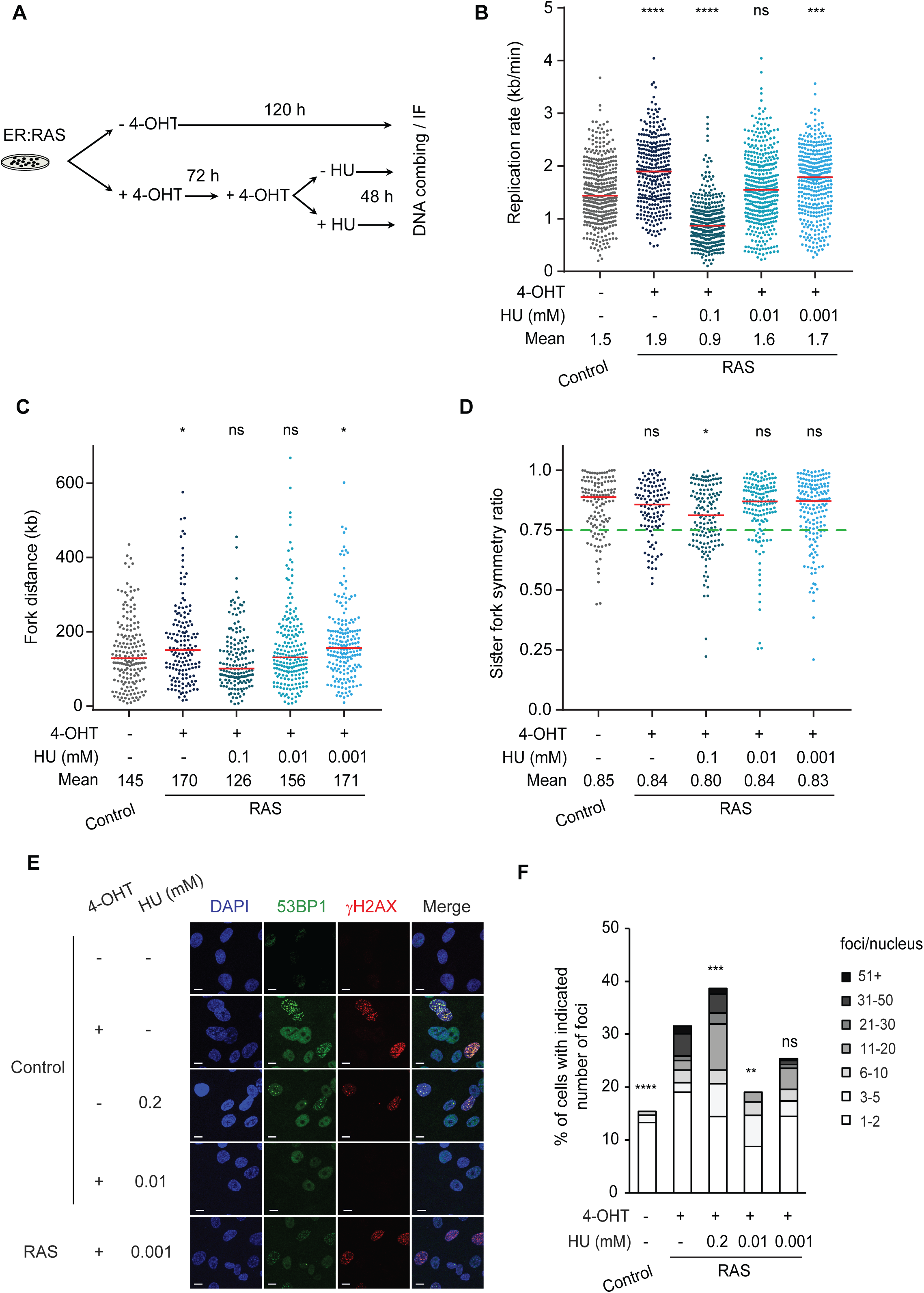
Mild replication inhibition restores normal replication dynamics and rescues DNA damage. (**A**) Scheme of the protocol. Cells infected with ER:RAS without 4-OHT treatment (Control) were cultured for 120h. Cells treated with 4-OHT (RAS) were cultured for 120h, with (+) or without (-) HU treatment for the last 48h, followed by DNA combing or immunofluorescence (IF). (**B-D**) DNA combing analysis of FSE-hTert cells with (+) or without (-) 4-OHT and HU treatments as indicated. Individual replication fork rates (kb/min); at least 300 fibers per condition were analyzed (**B**). Individual fork distances (kb); at least 150 forks per condition were analyzed (**C**). Individual sister fork symmetry ratios; at least 110 forks per condition were analyzed (**D**). Means are indicated; Red lines indicate medians. *P* values calculated compared to FSE-hTert cells without 4-OHT treatment (Control) by one-way ANOVA. Data are the summary of two independent experiments. (**D**) Dashed green line indicates asymmetry ratio threshold. (**E,F**) Co-localization of γH2AX (red) and 53BP1 (green) foci in ER:RAS FSE-hTert cells with (+) or without (-) 4-OHT and HU treatments as indicated. Representative images (**E**), quantification of the percent of cells with indicated number of foci per nucleus (**F**); number of analyzed nuclei from left to right: n = 286, 336, 194, 273, 276, respectively. Data are representative of two independent experiments with similar results. *P* values calculated compared to ER:RAS FSE-hTert cells treated with 4-OHT (+) but without HU (-) (RAS) by Mann Whitney rank-sum test. ns – non-significant; * *P* < 0.05, ** *P* < 0.01, *** *P* < 0.001, **** *P* < 0.0001. Scale bars, 10 μm.

Next, we investigated whether the replication rate restoration affected RAS-induced DNA damage. For this purpose, cells were treated with 0.01mM HU for 48 hours prior to analysis of DNA damage by the immunofluorescence detection of co-localized DNA damage markers γH2AX and 53BP1. Mild HU treatment in control cells did not lead to DNA damage induction (Figure S5A). However, as seen in Figure 3F, 0.01mM HU treatment led to a significant decrease in the number of RAS-induced foci compared with non-treated RAS-cells (Figures 3E and 3F). It is worth noting that the highest HU concentration (0.1mM) used caused a dramatic replication fork slowing in RAS expressing cells even when compared with control cells (Figure 3B). Accordingly, high HU treatment led to increased DNA damage formation compared with non-treated RAS-cells (Figures 3E and 3F). A considerably lower dose of HU (0.001mM) had a limited, non-significant effect on the replication rate (Figure 3B) and as expected had no significant effect on DNA damage formation compared with RAS expressing cells (Figures 3E and 3F). Altogether these results indicate that restoration of the accelerated replication rate dramatically reduced the DNA damage in RAS expressing cells.

We further investigated the effect of replication restoration on DNA damage by examining the effect of aphidicolin (APH), another replication inhibitor, which inhibits DNA polymerases α δ, ε and decreases fork progression in a dose-dependent manner (Cheng and Kuchta, 1993; Ikegami et al., 1978). RAS expressing cells were treated with relatively low APH concentrations for 48 hours prior to replication dynamics and DNA damage analyses. Flow cytometry analysis showed no significant change in cell cycle progression, indicating that like HU, APH treatment did not arrest cell proliferation (Figures S4C and S4D). Co-localization analysis of γH2AX and 53BP1 foci revealed that a low dose of 0.01 μM APH significantly decreased the number of foci in RAS expressing cells compared with non-treated RAS-cells (Figures S5C and S5D), while it had no effect on the level of the damage markers in control cells (Figure S5A). Furthermore, similar to the effect of various HU concentrations, a very low dose of APH (0.001 μM) did not have a significant effect on DNA damage formation in RAS-cells compared with non-treated RAS expressing cells (Figure S5D); by contrast, a high dose of M APH induced DNA damage formation (Figure S5D). Finally, we investigated whether the DNA damage rescue by the 0.01 μM APH treatment was associated with replication restoration. As expected, combing analysis showed restoration of the replication dynamics by the APH treatment (Figures S5E-S5G), altogether suggesting that in RAS cells accelerated replication generates DNA damage.

### Excess of Topoisomerase 1 levels causes an accelerated replication rate and DNA damage in RAS expressing cells

To explore the molecular mechanism/s underlying the accelerated replication in RAS-cells, we examined the differences in gene expression after mutated RAS activation. For this we performed RNA-seq analysis on control and RAS expressing cells at two time points, at two- and four-days post-RAS activation, when cells are proliferating. Principal component analysis showed that the expression profiles of RAS-cells clustered together and were distinguishable from the control cells (Figure S6A). After RAS activation >1,700 genes were differentially expressed, with an estimated false discovery rate (FDR) of < 5% and a fold change > 2-fold (Figure S6B). Gene Ontology (GO) annotation analysis of the upregulated genes following RAS activation (shared at both time points, 282 genes) showed enrichment for signaling and developmental processes (Figure S6C). Among the shared downregulated genes (568 genes) in RAS-cells, GO annotation analysis identified enrichment of anatomical and developmental processes (Figure S6C). DNA replication was not found among the GO annotations significantly enriched after RAS activation. Therefore, we next focused on individual DNA replication annotated genes (GO:0006260) to identify specific differentially expressed genes in RAS as compared with control cells, which could lead to dysregulation of the replication process. Previously, deregulation of origin firing factors such as CDC7, ORC1, MCM4, MCM6, Treslin and MTBP have been shown to lead to an increased replication rate in various organisms (Flach et al., 2014; Maya-Mendoza et al., 2018; Sedlackova et al., 2020; Zhong et al., 2013). However, our analysis showed no significant change in the expression level of any of these genes (Figure 4A and Supplementary Table 1), suggesting that in our system the accelerated replication rate was not the result of a decreased origin usage.

**Figure 4.**
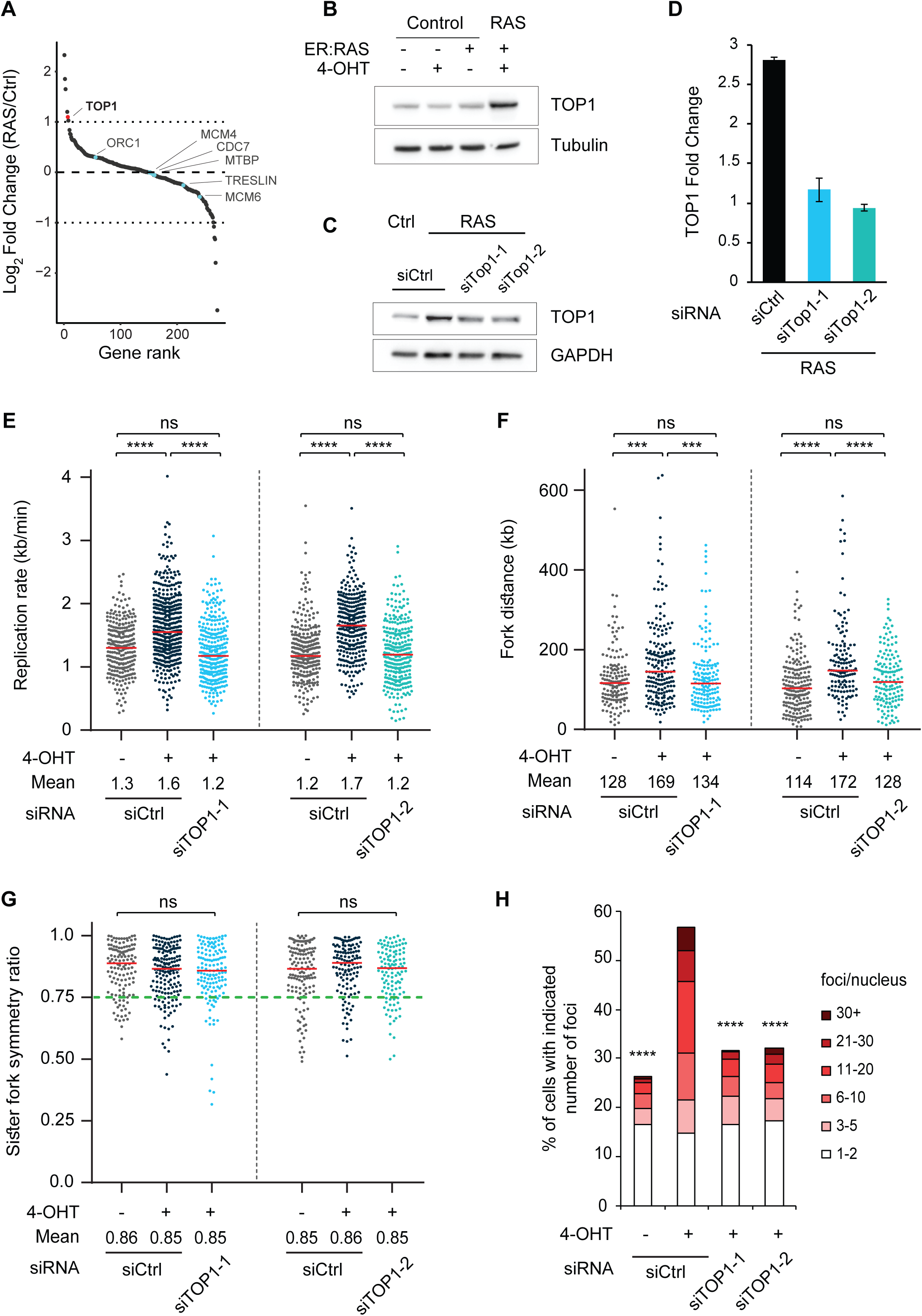
Increased TOP1 expression causes accelerated DNA replication and DNA damage. (**A**) Expressed DNA replication annotated genes (GO:0006260, n=258) ranked according to the RAS/Control fold change ratio. Light blue - genes previously associated with accelerated replication rate; red - TOP1. (**B**) Protein levels of TOP1 and Tubulin in FSE-hTert cells with (+) or without (-) ER:RAS infection and 4-OHT treatment as indicated. (**C**) Protein levels of TOP1 and GAPDH in ER:RAS infected FSE cells with (RAS) or without (Ctrl) 4-OHT treatment, and treated with two independent siRNAs against TOP1 (siTOP1-1 and siTOP1-2) or non-targeting siRNA (siCtrl), as indicated. (**D**) ER:RAS FSE hTert cells treated with 4-OHT (RAS) were treated with two independent siRNAs against TOP1 (siTOP1-1 and siTOP1-2) and non-targeting siRNA (siCtrl). Levels of TOP1 were measured by RT-qPCR and normalized to GAPDH. The values are averaged fold change (mean ± s.e.m, n = 2) relative to ER:RAS FSE hTert cells without 4-OHT treatment. (**E-G**) DNA combing analysis of ER:RAS FSE-hTert cells with (+) or without (-) 4-OHT and siRNA treatments as indicated. Individual replication fork rates (kb/min); at least 270 fibers per condition were analyzed (**E**). Individual fork distances (kb); at least 140 forks per condition were analyzed (**F**). Individual sister fork symmetry ratios; at least 125 forks per condition were analyzed (**G**). Means are indicated; Red lines indicate medians. *P* values calculated by one-way ANOVA. Data are the summary of two independent experiments. (**G**) Dashed green line indicates asymmetry ratio threshold. (**H**) Percent of cells with the indicated number of γH2AX foci in ER:RAS FSE-hTert cells with (+) or without (-) 4-OHT and siRNA treatments as indicated; number of analyzed nuclei from left to right: n = 2644, 1021, 2223, 3367, respectively. *P* values calculated compared to ER:RAS FSE-hTert cells treated with 4-OHT and siCtrl (RAS), Mann Whitney rank-sum test. Data are representative of three independent experiments. ns – non-significant; *** *P* < 0.001, **** *P* < 0.0001.

Analysis of replication annotated genes identified only five genes upregulated by at least two-fold following RAS activation: *E2F7*, *EREG*, *EGFR*, *HMGA1* and *TOP1* (Figure 4A and Supplementary Table 1). Interestingly, Topoisomerase 1 (TOP1) downregulation was reported to reduce replication fork rate and induce DNA damage (Promonet et al., 2020; Tuduri et al., 2009). TOP1 is an essential protein in mammalian cells that resolves the DNA torsional stress induced during replication and transcription (Pommier, 2006; Pommier et al., 2016). Therefore, we set to investigate the role of elevated TOP1 in regulation of the accelerated replication fork progression. First, we validated the increased level of TOP1 in RAS-cells by Western blot and RT-qPCR (Figures 4B and S7A). We then explored the relevance of TOP1 overexpression for cancers. Analysis of The Cancer Genome Atlas (TCGA) data sets revealed a two-fold increase in TOP1 in 9/33 cancer types, as found in our RAS-induced system (Figure S8A). We further identified in the TCGA datasets that TOP1 elevated expression positively correlates with the expression of RAS in 8/9 tested tumors (Figure S8B), further supporting a possible link between RAS activation and TOP1 overexpression.

To explore the effect of excess TOP1 levels on replication dynamics and DNA damage, we first restored normal TOP1 level in RAS-cells by moderate downregulation of TOP1 using low concentrations of two independent siRNAs (Figures 4C and 4D). We then examined the effect of restored TOP1 level on replication dynamics by DNA combing. The analysis showed that restoration of TOP1 to the normal expression level significantly reduced the replication rate in siTOP1-treated RAS-cells compared with control siRNA-treated RAS expressing cells (Figure 4E), which was indistinguishable from the rate of the control cells, indicating complete restoration of a normal replication rate (Figure 4E). TOP1 restoration also significantly reduced the fork distance in siTOP1-treated RAS- cells compared with control siRNA-treated RAS-cells (Figure 4F), indicating restoration of the replication dynamics. Finally, TOP1 restoration did not induce a significant increase in sister fork asymmetry, indicating that the mild siRNA downregulation did not induce fork stalling in RAS-cells (Figure 4G). Downregulation of TOP1 level in control cells using the same low concentrations of siTOP1 led to significant reduction in the replication fork rate and fork distance compared with treatment with a control siRNA (Figures S7B-S7D), in agreement of previous results (Promonet et al., 2020; Tuduri et al., 2009). Furthermore, downregulation of TOP1 in control cells significantly reduced the sister forks symmetry ratio (Figure S7E), suggesting that deficit in TOP1 leads to classical replication stress, whereas excess TOP1 drives a non-canonical form of replication stress characterized by fork acceleration.

We further investigated the effect of TOP1 restoration on RAS-induced DNA damage. For this we treated RAS-cells with siRNAs against TOP1, restoring normal expression level, and analyzed the DNA damage by immunofluorescence detection of γH2AX foci. The results showed a significant decrease in γH2AX levels following TOP1 downregulation in RAS-cells, compared with RAS-cells treated with control siRNA (Figure 4H). This indicates that restoration of TOP1 to the normal level of control cells, rescues the DNA damage formation. Thus, indicating that RAS-induced elevated level of TOP1 underlies the molecular mechanism of aberrant accelerated replication rate and genomic instability. Furthermore, downregulation of the TOP1 level in control cells, which led to a significant reduction in the replication rate, increased the level of DNA damage (Figure S7F), in agreement with previously published data (Promonet et al., 2020; Tuduri et al., 2009). Altogether these results indicate that regulated expression level of TOP1 is crucial since both increased and decreased TOP1 levels are deleterious to cells.

### Reduced TOP1-dependent R-loops promote accelerated replication and DNA damage

To test our hypothesis that excess TOP1 accelerates replication rate leading to DNA damage formation, we overexpressed TOP1 in HEK 293 cell (Figure 5A) and analyzed its effect on replication dynamics using DNA combing. TOP1 overexpression significantly increased the mean replication rate compared with control cells (Figure 5B), and accordingly increased the fork distance (Figure 5C). Fork symmetry analysis revealed no increase in asymmetric fork progression following TOP1 overexpression (Figure 5D), indicating that excess TOP1 expression accelerated DNA replication and did not cause fork stalling. Next, we measured DNA damage levels in TOP1 overexpressing cells by immunofluorescence analysis of the DNA damage marker 53BP1. The analysis showed an increase in the mean number of foci following TOP1 overexpression compared with control cells (Figure 5E), supporting our findings that TOP1-dependent accelerated replication leads to DNA damage formation.

**Figure 5.**
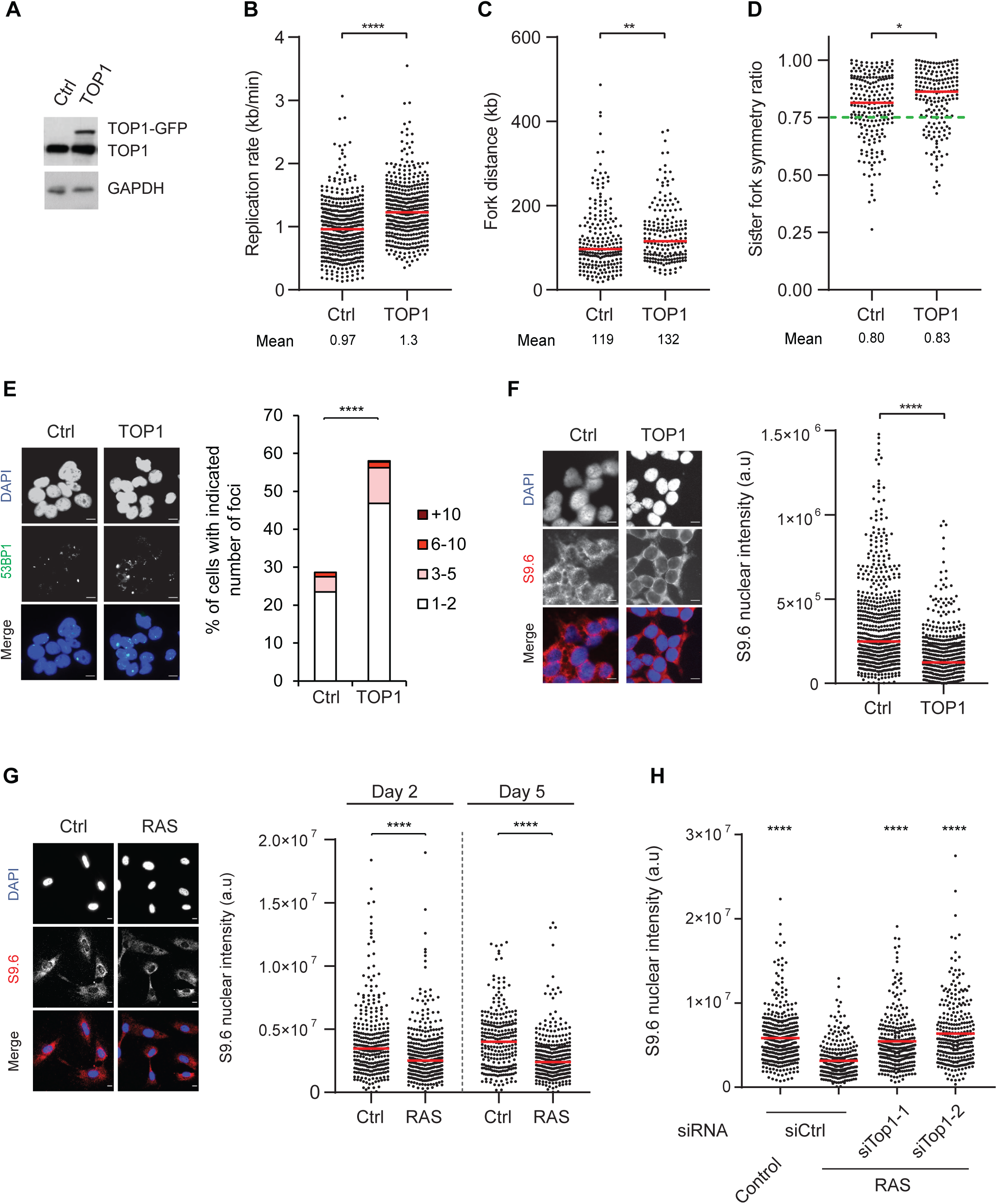
TOP1 overexpression reduces R-loop level leading to accelerated DNA replication and damage. (**A**) Protein levels of endogenous TOP1 (TOP1), ectopic TOP1 (TOP1-GFP) and GAPDH in HEK-293 cells transfected with a control GFP vector (Ctrl) or with TOP1-GFP (TOP1), as indicated. (**B**-**D**) DNA combing analysis of HEK-293 cells as indicated in **A**. Individual replication fork rates (kb/min); at least 400 fibers per condition were analyzed (**B**). Individual fork distances (kb); at least 180 forks per condition were analyzed (**C).** Individual sister fork symmetry ratios; at least 180 forks per condition were analyzed (**D**). Means are indicated; Red lines indicate medians. *P* values calculated by Mann Whitney rank-sum test. Data are the summary of two independent experiments. Dashed green line indicates asymmetry ratio threshold (**D**). (**E**) 53BP1 foci in HEK-293 cells as indicated in **A**. Left: Representative images of 53BP1 (green) foci and DAPI (blue) staining. Right: Quantification of the percent of cell with indicated number of 53BP1 foci. At least 300 nuclei were analyzed. Data are representative of two independent experiments. (**F**) RNA-DNA hybrids in HEK-293 cells as indicated in **A**. Left: Representative images of RNA-DNA specific antibody (S9.6, red) and DAPI (blue) staining. Right: Dot plot of mean nuclear fluorescence intensity of RNA-DNA hybrid specific antibody (S9.6) in individual nuclei. At least 600 nuclei were analyzed. Data are summary of two independent experiments. (**G**) RNA-DNA hybrids in ER:RAS FSE hTert cells with (RAS) or without (Ctrl) 4-OHT treatment. Left: representative images of RNA-DNA specific antibody (S9.6, red) and DAPI (blue) staining. Right: Dot plots of mean nuclear fluorescence intensity of RNA-DNA hybrids (S9.6 antibody) in individual nuclei in control (Ctrl) and RAS expressing cells for 2 or 5 days. (**H**) Dot plot of mean nuclear fluorescence intensity of RNA-DNA hybrid specific antibody (S9.6) in individual nuclei in ER:RAS FSE hTert cells with (RAS) or without (Ctrl) 4-OHT treatment, and siRNA treatments as indicated. * *P* < 0.05; ** *P* < 0.01; **** *P* < 0.0001. Scale bars, 10 μm.

We hypothesize that elevated TOP1 could promote accelerated replication by reducing the abundance of R-loops and therefore decrease potential obstacles to the replication machinery. R-loops are RNA-DNA hybrids formed during transcription and their aberrant accumulation may stall replication fork progression and thus lead to DNA damage (Santos-Pereira and Aguilera, 2015). Negative supercoiling, formed behind the transcribing RNA polymerase, promotes DNA strand separation which in turn increases the possibility of the nascent transcript to anneal to the DNA, and form an R-loop (Santos-Pereira and Aguilera, 2015). TOP1 relieves the negative supercoiled DNA behind the transcription machinery and thus prevents R-loop formation (Santos-Pereira and Aguilera, 2015). Indeed, TOP1 inhibition or down-regulation increases R-loop level (Promonet et al., 2020; Tuduri et al., 2009). Therefore, we investigated the effect of elevated TOP1 on R-loop accumulation. Immunofluorescence analysis of RNA-DNA hybrids indeed showed a significant R-loop decrease in TOP1 overexpressing cells compared with control cells (Figure 5F), implying that reduced R-loop levels by increase TOP1 levels may promote accelerated replication rate.

Next, we explored the effect of excess TOP1 on R-loop levels in RAS expressing cells. Analysis of RNA-DNA hybrid levels showed a significant decrease in R-loops in RAS expressing cells for 2 and 5 days compared with control cells (Figure 5G). We then analyzed the effect of TOP1 restoration in RAS-cells on R-loops, using two independent siRNAs against TOP1. The results showed that the siTOP1 treatment increased R-loop levels in RAS-cells compared with control siRNA-treated RAS-cells (Figure 5H). Notably, TOP1 restoration by siRNA increased R-loop levels back to normal (Figure 5H). These results indicate that restoration of TOP1 level in RAS-cells restores R-loop levels and rescues the accelerated replication rate and DNA damage. Similarly, mild replication inhibition by HU/APH, which restored normal replication dynamics (Figures 3 and S5), also increased R-loop levels back to normal, without affecting TOP1 level (Figure S9). Altogether, these results suggest that R-loop restoration rescues the perturbed DNA replication.

Finally, we explored the direct effect of R-loop suppression on replication dynamics. For this, we overexpressed RNaseH1, which degrades R-loops, in HEK 293 cells (Figure 6A). As expected RNaseH1 overexpression reduced the level of R-loops compared with control cells (Figure 6B). RNaseH1 overexpression significantly accelerated the replication rate compared with control cells (Figure 6C), and accordingly increased the mean fork distance (Figure 6D). Fork symmetry analysis revealed no change following RNaseH1 overexpression (Figure 6E). Lastly, immunofluorescence analysis of the DNA damage marker 53BP1, showed an increase in the mean number of foci following RNaseH1 overexpression compared with control cells (Figure 6F). Taken together, the results indicate that suppressed R-loop levels promote replication fork acceleration and lead to the formation of DNA damage.

**Figure 6.**
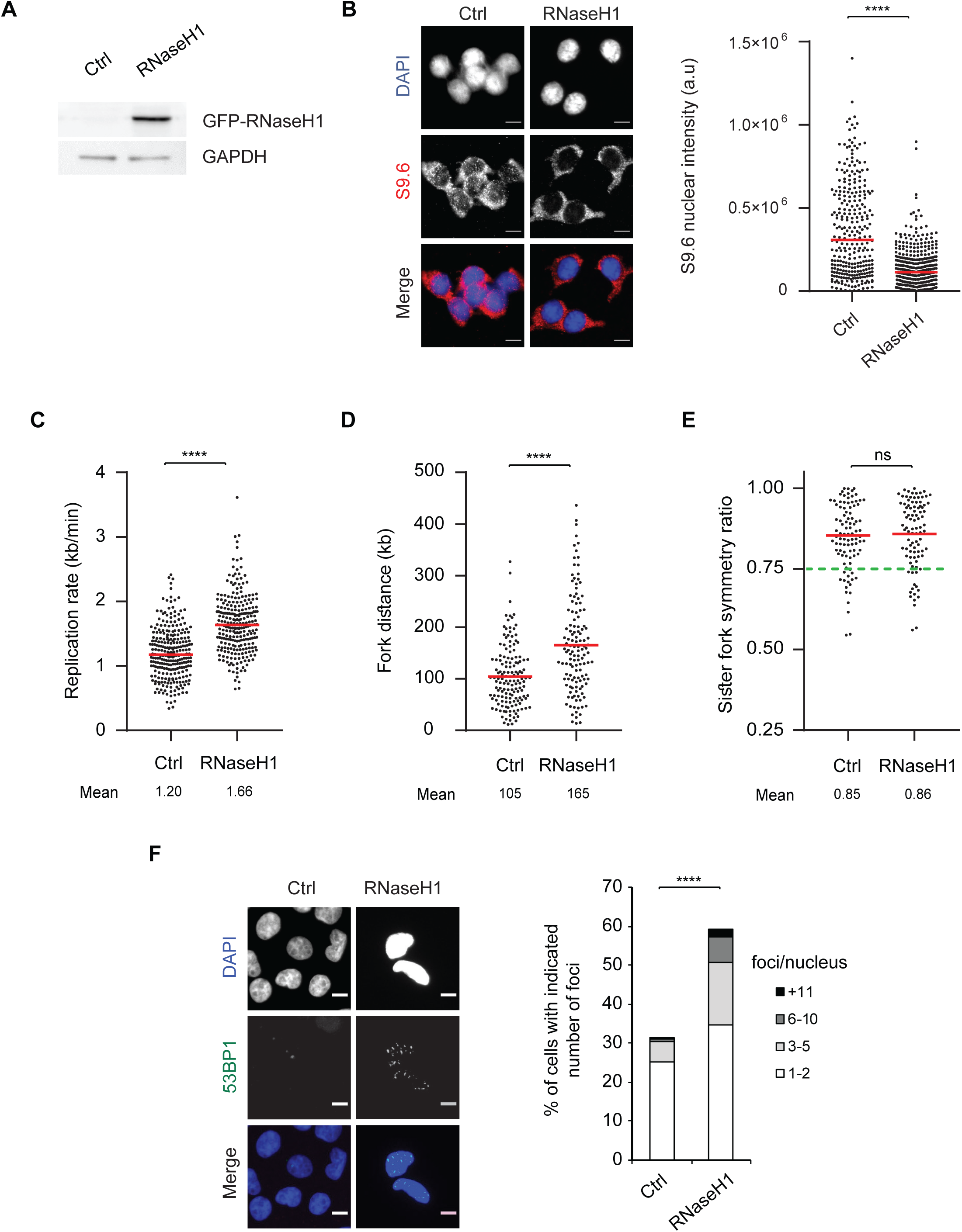
Reduced R-loops causes accelerated DNA replication rate and DNA damage. (**A**) Protein levels of GFP-RNaseH1 and GAPDH in HEK-293 cells transfected with a control GFP plasmid (Ctrl) or with GFP-RNaseH1 plasmid (RNaseH1). (**B**) RNA-DNA hybrids in HEK-293 cells transfected with a control GFP plasmid (Ctrl) or with a GFP-RNaseH1 plasmid (RNaseH1). Left: Representative images of RNA-DNA specific antibody (S9.6, red) and DAPI (blue) staining. Right: Dot plot of mean nuclear fluorescence intensity of RNA-DNA hybrid specific antibody (S9.6) in individual nuclei of Ctrl (n = 291) and RNaseH1 (n = 439). Data are the summary of two independent experiments. (**C-E**) DNA combing analysis of HEK-293 cells transfected with a control GFP plasmid (Ctrl) or with GFP-RNaseH1 plasmid (RNaseH1). Individual replication fork rates (kb/min) in Ctrl and RNaseH1 cells; at least 250 fibers per condition were analyzed (**C**). Individual fork distances (kb) in Ctrl and RNaseH1 cells; at least 130 fibers per condition were analyzed (**D**). Individual sister fork symmetry ratios in Ctrl and RNaseH1 cells; at least 90 fibers per condition were analyzed (**E**). Means are indicated; Red lines indicate medians. *P* values calculated by Mann Whitney rank-sum test. Data are the summary of two independent experiment. Dashed green line indicates asymmetry ratio threshold (**E**). (**F**) 53BP1 foci in HEK-293 cells transfected with a control GFP plasmid (Ctrl) or with GFP-RNaseH1 plasmid (RNaseH1). Left: Representative images of 53BP1 (green) foci and DAPI (blue) staining. Right: quantification of the percent of cell with indicated number of 53BP1 foci in Ctrl (n = 1120) and RNaseH1 (n = 1111). Data are the summary of two independent experiments. ns – non-significant; **** *P* < 0.0001. Scale bars, 10 μm.

## Discussion

In this study we highlight the importance of TOP1 equilibrium in the regulation of R-loop homeostasis to ensure faithful DNA replication and genome integrity. We show that an increase in TOP1 expression leads to a reduction in R-loop levels promoting replication fork acceleration resulting in DNA damage (Figure S10).

TOP1 is an essential protein for both DNA replication and transcription, resolving topological stress generated by DNA strand supercoiling (Pommier et al., 2016). During DNA replication TOP1 resolves positive supercoiling, formed ahead of the replication fork, allowing the persistent progression of the replisome (Pommier et al., 2016). During transcription TOP1 resolves negative supercoiled DNA, formed behind the RNA polymerase. Such negative supercoiling facilitates the hybridization of the nascent transcript to the template DNA generating an RNA-DNA hybrid (R-loop) (Pommier et al., 2016). R-loops have a role in regulating gene expression and mediating transcription termination (Promonet et al., 2020; Santos-Pereira and Aguilera, 2015). However, persistent R-loop accumulation is a major threat to genome integrity, as it introduces potential barriers for the replicating fork progression (Bhatia et al., 2014; Costantino and Koshland, 2018; Kotsantis et al., 2016; Parajuli et al., 2017; Santos-Pereira and Aguilera, 2015; Tuduri et al., 2009). Indeed, TOP1 downregulation or inhibition increases R-loop levels and slows the replication rate (Promonet et al., 2020; Tuduri et al., 2009; Xu and Her, 2015). Moreover, TOP1 knockout mice are embryonically lethal (Morham et al., 1996). In contrast to TOP1 deficiency, little is known about the effect of excess TOP1. Transient TOP1 overexpression was associated with DNA damage (Humbert et al., 2009), and elevated levels of TOP1 are found in several cancers (Figure S8) (Grunnet et al., 2015; Lee et al., 2015; Rømer et al., 2012). However, whether surplus TOP1 affects replication dynamics was unexplored. Here we show that increased expression of TOP1 reduces the levels of R-loops, accelerates DNA replication and generates DNA damage (Figures 5 and 6). Mild TOP1 downregulation using siRNA restored TOP1 level in RAS expressing cells back to normal expression level of control cells. Following TOP1 restoration R-loop levels were elevated back to normal, replication rate was restored, and the damage was rescued (Figure 4 and 5). We further show that R-loop depletion by RNaseH1 overexpression also increases the rate of replication and induces DNA damage (Figure 6), indicating that TOP1-dependent R-loop deficiency underlies replication fork acceleration and DNA damage. Together, our results highlight the importance of a regulated expression of TOP1 for R-loop equilibrium, ensuring faithful DNA replication and genome integrity, as both excess and insufficient levels perturb DNA replication and lead to DNA damage. It should be noted that we cannot exclude the possibility that TOP1-dependent DNA replication acceleration may also result, in a non-mutually exclusive manner, from its role in resolving positive supercoiled DNA ahead of the moving replication machinery. Future investigation may shed light on this role.

Replication stress is mainly characterized by fork slowing and stalling generating DNA damage (Zeman and Cimprich, 2014). While this is largely true today, an increasing body of evidence accumulated in recent years has unveiled the diverse nature of replication stress, as other forms of perturbed replication were reported. Among the emerging non-canonical forms of replication stress is replication fork acceleration. Previous studies found accelerated replication rate progression following downregulation of mRNA biogenesis genes, including THOC1, PCID2 and GANP (Bhatia et al., 2014; Domínguez-Sánchez et al., 2011), and origin licensing and firing: ORC1, CDC7, CDC6 and MCM4 (Sedlackova et al., 2020; Zhong et al., 2013). Interestingly however, no change in the expression levels of mRNA biogenesis or origin associated genes was found in our RAS-cells (Supplementary Table 1 and Figure 4A), supporting that various mechanisms underlie accelerated replication rate progression.

Recently, accelerated replication was reported following dysregulation of cancer associated genes, including overexpression of the oncogene Spi-1 (Rimmele et al., 2010), interferon-stimulated gene 15, ISG15 (Raso et al., 2020), and following exposure to the developmental mitogen sonic hedgehog, Shh (Tamayo-Orrego et al., 2020). Moreover, inhibition of poly(ADP-ribose) polymerase (PARP), a common anti-tumor agent that facilitates repair of DNA damage, accelerated the replication rate and promoted genomic instability (Maya-Mendoza et al., 2018; Sugimura et al., 2008). Interestingly, inhibition of PARP, delocalizes TOP1 from the nucleolus to the nucleoplasm (Das et al., 2016) where replication of the bulk of the genome takes place. Thus, higher levels of TOP1 in the nucleoplasm may contribute to the observed accelerated replication rate after PARP inhibition.

Most studies reporting accelerated DNA replication also observed an increase in DNA damage. However, whether the accelerated replication directly causes damage remained unknown. Here, we addressed this question by mild inhibition of replication fork progression using low doses of HU/APH treatment, restoring normal replication dynamics in RAS expressing cells (Figures 3 and S5). Following replication restoration, the DNA damage was significantly reduced, indicating that accelerated replication drives DNA damage formation. In order to test the possibility that the low HU/APH treatments activated the DDR, which might have contributed to the rescue of damage, we treated control cells with corresponding low HU/APH concentrations and found no DDR activation (Figure S5A), in agreement with a previous report (Koundrioukoff et al., 2013). This indicates that the DNA damage rescue by mild replication inhibition is due to replication restoration. Altogether, our results indicate that accelerated replication directly generates DNA damage.

The damage induced by accelerated replication is mechanistically distinct from damage induced by decelerated replication rate. Previous studies of oncogene-induced replication stress driving genomic instability showed that slow replication-induced DNA damage resulted from either insufficient levels of dNTPs (Bester et al., 2011) or from an increase in replication-transcription conflicts manifested by increased levels of R-loops (Jones et al., 2013). However, our results show that accelerated replication-induced damage is not the result of a deficiency in dNTP levels, as supplementation of dNTPs did not affect the damage levels in RAS expressing cells (Figure S3). Furthermore, accelerated replication-induced damage could not result from an increased level of R-loops, since we observed reduced levels of R-loops in TOP1 overexpressing cells and RAS-expressing cells (Figures 5F and 5G). While exploring the mechanism underlying accelerated replication-induced DNA damage is beyond the scope of this study, it is reasonable to speculate that the increased fork progression impairs DNA polymerase fidelity, leading to DNA damage formation by increased mutagenesis (Kunkel, 2004).

In previous studies in which mutated RAS was overexpressed, various effects on replication dynamics were found. Mendoza et al., reported a small yet a significant increase in replication rate at the hyperproliferative phase, prior to senescence onset. However, this was followed by a decrease in replication rate and an increase in DNA damage levels (Maya-Mendoza et al., 2015). In another study, Di Micco et al., overexpressed RAS in CHK2 deficient cells, which prevented RAS-induced senescence. The effect of RAS overexpression on the replication dynamics was analyzed by measuring inter-origin distance (Di Micco et al., 2006). The results showed a shorter inter-origin distance indicating an increased origin firing. Kotsantis et al., reported that RAS overexpression resulted in slow replication and formation of DNA damage. The authors explored the mechanism underlying the slow replication, and showed that RAS upregulated the general transcription factor TATA-box binding protein (TBP) at the hyperproliferative phase, which led to an increase in RNA synthesis, accumulation of R-loops and slow rate and DNA damage (Kotsantis et al., 2016). In our study, however, RAS overexpression did not change TBP expression (Supplementary Table 1). Furthermore, RAS expression in our study reduced R-loop levels due to TOP1 overexpression. Different factors may underlie the various effects of RAS overexpression on the replication dynamics in these studies. One possible factor is a different RAS overexpression level, since a dose-dependent effect of RAS expression on cell proliferation and senescence was previously shown (Sarkisian et al., 2007), which may reflect differences in replication dynamics and DNA damage. Another possible factor is the use of different cellular systems, which may have a different genetic background. Furthermore, it is well established that there is accumulation of genetic changes during cell culturing of cancer cell lines, leading to variations in gene expression patterns which may differently affect the replication dynamics and DNA damage (Ben-David et al., 2018; Kim et al., 2017; Weissbein et al., 2014). Altogether, the plasticity of RAS-induced replication stress found in different studies suggests that the genetic background of oncogenic cells may affect the phenotype and hence should be reflected in the treatment for cancer patients.

Altogether, our results indicates that TOP1 is a crucial factor for genome integrity, by tightly regulating replication dynamics. This further highlights the importance of TOP1- dependent R-loops homeostasis in replication regulation, as unbalanced levels impede replication dynamics and promote genomic instability. These results are highly important for understanding early events leading to cancer development, as different mechanisms may underlie the oncogene-induced replication perturbation driving genomic instability. This sheds light on the complex nature of oncogene-induced replication stress and suggests it should be taken into consideration when replication stress is considered as a therapeutic tool for cancer.

## Methods

### Cell Culture

Human foreskin fibroblasts, FSE-hTert cells, lung fibroblasts, WI38-hTert, and HEK-293 cells were grown in DMEM supplemented with 10% fetal bovine serum, 100,000 U l^-1^ penicillin and 100 mg l^-1^ streptomycin. ER:RAS activation was induced by supplementing the growth media with 200 μM of 4-hydroxytamoxifen (4-OHT). Nucleoside supplementation was achieved by supplementing the growth media with 50 µM of A, U, C and G each for 48 h prior to fixation. Aphidicolin and hydroxyurea treatments were performed in growth media with indicated concentrations for 48 h prior to fixation.

### Plasmids

For ER:RAS infection, Phoenix retroviral packaging cells were transiently transfected with ER:RAS pLNC vector plasmids (kindly provided by Dr. J.C Acosta). Cells were infected three times with the Phoenix cell supernatant, containing replication-defective viruses. Infected FSE and WI38 cells were selected using 400 μg ml^-1^ G418 for the next 10 days. For TOP1-GFP transfection, HEK-293 cells were transiently transfected with pEGFP-TOP1 (kindly provided by Dr. Tasuku Honjo) or with a control GFP pBabe vector. Transfected cells were FACS sorted 24 h post transfection and 24 h later GFP positive cells were analyzed. For GFP-RNaseH1 transfection, HEK-293 cells were transiently transfected with pEGFP-RNaseH1 vector (kindly provided by Dr. Robert Crouch) or with a control GFP pBabe vector. Transfected cells were FACS sorted 24 h post transfection and 48 h later GFP positive cells were analyzed.

### Replication dynamics using DNA combing

Molecular combing is a process whereby single DNA molecules (hundreds of Kbs) are stretched on a silanized glass surface (COV-002, Genomic vision) (Bensimon et al., 1994). In general, unsynchronized cells were labeled for 30 min by medium containing 100 μ of the thymidine analog iododeoxyuridine (IdU). At the end of the first labeling period, the cells were washed twice with a warm medium and pulse labeled once more for 30 min with a medium containing 100 μM chlorodepxyuridine (CldU) and then washed with cold PBS and harvested. Genomic DNA was extracted using Fiber-Prep kit (EXTR-001, Genomic Vision), combed using the Fiber-Comb (MSC-001, Genomic Vision) and analyzed as previously described (Herrick and Bensimon, 1999). The primary antibody for fluorescence detection of IdU was mouse anti-BrdU (Becton Dickinson), and the secondary antibody was goat anti-mouse Alexa Fluor 488 (Invitrogen). The primary antibody for fluorescence detection of CldU was rat anti-CldU (Novus Biologicals). The secondary antibody was goat anti-rat Alexa Fluor 594 (Invitrogen). The primary antibody for fluorescence detection of ssDNA was mouse anti-ssDNA (Millipore). The secondary antibody was donkey anti-mouse Alexa Fluor 647 (Invitrogen). The length of the replication signals and the fork distances were measured in micrometers and converted to kilo bases according to a constant and sequence-independent stretching factor (1μm= 2kb), as previously reported (Herrick and Bensimon, 1999). Images were analyzed double blindly using Fiji (Schindelin et al., 2012).

### Immunofluorescence staining

Cells were fixed in 4% formaldehyde/PBS for 10 min, permeabilized with 0.5% Triton/PBS for 10 min, and blocked with 10% fetal bovine serum/PBS for 1-3 h. The primary antibodies used were mouse anti-phosphorylated H2AX (Upstate Biotechnology, 1:100), rabbit anti-53BP1 (Bethyl Laborartories, 1:100), mouse anti-cyclin A2 (Abcam, 1:100) and mouse anti-BrdU (Becton Dickinson, 1:25). For RNA-DNA hybrid detection, cells were washed with cold PBS and fixed with 100% ice-cold methanol for 7 min and incubated overnight in blocking solution 3% BSA/PBS. S9.6 antibody (kindly provided by Dr. Rachel Eiges) was used, 1:500. Secondary antibodies added were anti–mouse Alexa Fluor 488 (Invitrogen), anti-rabbit Alexa Fluor 488 (Invitrogen), anti-mouse Alexa Fluor 555 (Invitrogen). DNA was counterstained with mounting medium for fluorescence with DAPI (Vector Laboratories). For focus information analysis images were taken with the FV-1200 confocal microscope (Olympus, Japan), with a 60X/1.42 oil immersion objective. Multiple dyes sequential scanning mode was used in order to avoid emission bleed-through. For focus and fluorescent intensity analysis the Hermes WiScan system (Idea Bio-Medical, Israel) was used. All images were analyzed double blindly using Fiji (Schindelin et al., 2012).

### Metaphase chromosome preparation and fragile site analysis

Cells were treated with 100 ng ml^-1^ colcemid (Invitrogen) for 15–40 min, collected by trypsinization, treated with hypotonic solution at 37 °C for 30 min and fixed with multiple changes of methanol:acetic acid 3:1. Fixed cells were kept at -20°c until analysis. For analysis of total gaps and breaks chromosomes were stained with propidium-iodide and analyzed double blindly using Fiji (Schindelin et al., 2012).

### Western blot analysis

8-12% polyacrylamide gels were used for protein separation and detection. The gels were transferred to a nitrocellulose membrane, and antibody hybridization and chemiluminescence (ECL) were performed according to standard procedures. The primary antibodies used in these analyses were rabbit anti-H-RAS (Santa Cruz, 1:1,000), mouse anti- CHK1 (Cell signaling, 1:500), rabbit anti-phosphorylated CHK1 (Cell Signaling, 1:200), rabbit anti-GAPDH (Cell Signaling, 1:1,000), mouse anti-β-catenin (BD-Biosciences, 1:2,500), mouse anti-Tubulin (Sigma, 1:50,000), rabbit anti-TOP1 (Abcam, 1:10,000)), and rabbit anti-RNaseH1 (Abcam, 1:1,000). HRP-conjugated anti-rabbit and anti-mouse secondary antibodies was obtained from Jackson Immunoresearch Laboratories (711-035-152, 1:5,000).

### Population doublings

Cells were grown in media as indicated in the ‘Cell Culture’ section. Initial seeding concentration of cells was 10,000 per well. Cells were trypsinized and counted. Population doublings (PD) was measured according to the following formula:

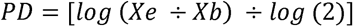

Xb is the number of cells at the beginning of incubation and Xe is the number of cell at the final count (Greenwood et al., 2004).

### RNA sequencing analysis

Sequencing libraries were prepared using the Illumina TruSeq mRNA kit, and sequenced (60 bp, single reads) on a single lane of Illumina HiSeq 2500 V4 instrument, to a depth of ∼27 million reads per sample. Reads were aligned to the hg19 genome (UCSC, downloaded from iGenomes) using TopHat (v2.0.10) (Kim et al., 2013). HTSeq-count (version 0.6.1p1) (Anders et al., 2015) was used to count reads on gene exons (UCSC Annotation from March 9, 2012). Differential expression analysis was performed using DESeq2 (1.6.3) (Love et al., 2014) with betaPrior set to False. Gene set enrichment analysis was performed using WebGestalt (Wang et al., 2013).

### Gene correlation analysis in GEPIA

The online database Gene Expression Profiling Interactive Analysis (GEPIA) (Tang et al., 2017) was used to explore TOP1 expression level in cancer. GEPIA is an interactive tool for analyzing RNA-seq expression data from 9,736 tumors and 8,587 normal samples from the TCGA and GTEx projects. GEPIA was used to generate TOP1 expression plot in normal and tumor samples and gene expression correlation analysis in tumor samples.

### RNA analysis

Total RNA was extracted using the RNeasy Mini Kit extraction kit (QIAGEN). RNA-less and reverse transcriptase-less reactions were used as controls. Complementary DNA (cDNA) synthesis was performed using the High Capacity cDNA Reverse Transcription kit (Applied Biosystems). Real-time PCR was subsequently performed in ABI 7500 using a Power SYBR green PCR master Mix (Applied Biosystems). The expression level was normalized to the transcript levels of GAPDH. Specific primers for these PCRs were designed using the Primer Express software: GAPDH: Fwd, TGAGCTTGACAAAGTGGTCG; Rev, GGCTCTCCAGAACATCATCC, POLR2A: Fwd, TGCGCA CCATCAAGAGAGTC; Rev, CTCCGTCACAGACATTCGCTT, TOP1: Fwd, CCCTGTACTTCATCGACAAGC; Rev, CCACAGTGTCCGCTGTTTC.

### siRNA

siRNA against TOP1 (TOP1-1: 5’-GCACAUCAAUCUACACCCA-3’ and TOP1-2: 5’-CGAAGAAGGUAGUAGAGUC-3’) and a control, non-targeting siRNA (Ctrl: 5’-UGGUUUACAUGUCGACUAA-3’) were purchased from Dharmacon. Cells were transfected with 40nM control siRNA and 20nM siRNA against TOP1, using oligofectamine (Thermo-Fisher). Cells were analyzed 48 hours after transfection.

### Cell cycle analysis

Cells were harvested and the pellet resuspended in 0.5ml cold PBS and fixed in 4.5ml 100% chilled methanol and kept at -20°C. Prior to FACS analysis, methanol residues were washed and cells were resuspended in PBS containing 0.2µg/µl RNase for 30 min. Cells were stained with 50µg/ml propidium iodide and the DNA content was analyzed by flow cytometry (BD FACSAria III).

### Statistical analysis

All data analysis was performed using Excel, GraphPad Prism 8.3.0 for Windows, GraphPad Software, La Jolla California USA (www.graphpad.com) or R project for Statistical Computing (http://www.r-project.com). For comparisons of replication dynamics (relevant to Figures 1,3,4,5,6 and S2,5,7), Immunofluorescence staining (relevant to Figures 2,3,4,5,6 and Figures S1,3,5,7,9), metaphase spreads analyses (relevant to Figure 2) and SA-β gal activity (relevant to Figure S1) one-way ANOVA and Mann-Whitney rank sum test were performed, as indicated. Numbers of repeats are indicated in the figure legends.

### Data availability

The authors declare that all the data supporting the findings of this study are available within the article and its Supplementary Information files and from the corresponding authors upon reasonable request. All sequencing files and processed count matrices were deposited in Gene Expression Omnibus (GEO) and will be available upon publication.

## Acknowledgements

This research was supported by grants from the Israel Science Foundation (grant No. 176/11), the Israeli Centers of Research Excellence (I-CORE), Gene Regulation in Complex Human Disease, Center No 41/11/”, and by the ISF-NSFC joint program (grant No. 2535/16). The authors thank Dr. Naomi Melamed-Book for her assistants in confocal microscopy. The authors thank the Mantoux Bioinformatics institute of the Nancy and Stephen Grand Israel National Center for Personalized Medicine, Weizmann Institute of Science, for assistant in deep sequencing and bioinformatics analysis. The authors thank the members of the Kerem lab for thoughtful discussions and advice.

## Author contributions

D.S contributed to conception and design, performed experiments, collection and assembly of data, data analysis and interpretation and manuscript writing; A.S performed and contributed to the experimental analyses, data interpretation and graphical abstract; Y.S.O performed and contributed to the experimental analyses, data interpretation and manuscript writing; B.K contributed to conception and design, financial support, data analysis and interpretation, manuscript writing and final approval of the manuscript.

## Supplementary legends

**Supplementary Figure 1. Related to Figure 1. RAS expression induces senescence.** (**A**) Quantification of population doublings at the indicated time points in: FSE-hTert cells (hTert, n = 4); FSE-hTert with 4-OHT treatment (hTert + 4-OHT, n = 2); FSE-hTert with ER:RAS infection but without 4-OHT treatment (ER:RAS, n = 4); FSE-hTert with ER:RAS infection and 4-OHT treatment (ER:RAS + 4-OHT, n = 4). n represents the number of independent experiments; data are mean ± s.e.m. (**B**) Representative images of IdU (green) and DAPI staining (blue) in RAS expressing cells at the indicated time points. (**C**) Quantification of the percent of control (Ctrl) and RAS nuclei positive for IdU incorporation at the indicated time points (days post RAS activation). Data are mean ± s.e.m from two independent experiments. (**D**) Representative images of senescence associated β-gal activity, positive cells are stained blue. (**E**) Quantification of senescence associated β-gal activity in FSE-hTert cells with (+) or without (-) ER:RAS infection and 4-OHT treatment at the indicated time points (days). Data are the summary of two independent experiments. *P* values calculated compared to FSE-hTert cells without ER:RAS infection nor 4-OHT treatment. Mann Whitney rank-sum test, ns - nonsignificant; **** *P* < 0.0001. Scale bars, 20 μm.

**Supplementary Figure 2. Related to Figure 1. RAS expression leads to increased replication rate and fork distance in WI38 cells.** (A) Protein levels of HRAS and GAPDH in WI38-hTert cells with (+) or without (-) ER:RAS infection and 4-OHT treatment as indicated. (B) Dot plots of individual replication fork rates (kb/min) in WI38-hTert infected with ER:RAS without (Control) or with (RAS) 4-OHT treatment for 5 days; at least 600 fibers per condition were analyzed. (C) Dot plots of individual fork distances (kb) in Control and RAS cells; at least 260 fibers per condition were analyzed. (D) Dot plots of sister fork symmetry ratios in Control and RAS cells; at least 230 fibers per condition were analyzed. Dashed green line indicate asymmetry ratio threshold. (B-D) Red lines indicate medians, means are indicated. (E) Scatter plots of the replication rates (kb/min) of right- and left-moving sister forks during IdU pulse. The center areas delimited with red lines contain sister forks with less than 25% difference. The percentages of symmetric forks are indicated at the top left corner of the plots. (B-E) Data are the summary of two independent experiments, Mann-Whitney rank-sum test, ns - nonsignificant; * *P* < 0.05; **** *P* < 0.0001.

**Supplementary Figure 3. Related to Figure 2. DNA damage induced by RAS expression is not a result of nucleotide insufficiency.** (A) Representative images of 53BP1 foci (green) in cyclin A (red) negative cells and DAPI staining (blue) in FSE-hTert cells infected with ER:RAS and treated with 4-OHT for 5 days with (+) or without (-) nucleoside supplementation (dNTPs). (B) Quantification of the mean 53BP1 foci in cyclin A negative ER:RAS FSE-hTert cells with 4- OHT treatment for 5 days, with exogenously supplementation of nucleosides (n = 273) or without (n = 293). Results are the summary of two independent experiments. Means ± s.e.m are shown. Mann-Whitney rank-sum test, ns - nonsignificant. Scale bars, 20 μm.

**Supplementary Figure 4. Related to Figure 3. Mild replication inhibition does not affect cell cycle progression.** (**A**) Flow cytometry analysis of ER:RAS FSE-hTet cells treated with 4- OHT and HU as indicated. (**B**) Quantification of data presented in **A**. (**C**) Flow cytometry analysis of ER:RAS FSE-hTet cells treated with 4-OHT and APH as indicated. (**D)** Quantification of data presented in **C**. Data are representative of two independent experiments.

**Supplementary Figure 5. Related to Figure 3. Mild replication inhibition restores normal replication dynamics and rescues DNA damage.** (A) Percent of cells with the indicated number of co-localized γH2AX and 53BP1 foci in ER:RAS infected FSE-hTert cells without 4-OHT treatment (control), treated with HU and APH as indicated. Cells treated with 0.2μM APH as a control, using a high APH concentration that generates DNA damage. Data are the summary of two independent experiments. (B) Scheme of the protocol. Cells infected with ER:RAS without 4-OHT treatment (control) were cultured for 120h. Cells treated with 4-OHT (RAS) were cultured for 120h, with (+) or without (-) aphidicolin (APH) treatment for the last 48h, followed by DNA combing or immunofluorescence (IF). (C,D) Co-localization of γH2AX (red) and 53BP1 (green) foci in ER:RAS FSE-hTert cells with (+) or without (-) 4-OHT and APH treatment at the indicated concentrations. Representative images (C), quantification of the percent of cells with indicated number of foci (D); number of analyzed nuclei from left to right: n = 142, 137, 142, 164, 141, respectively. Data are representative of two independent experiments with similar results. *P* values calculated compared to ER:RAS FSE-hTert cells treated with 4-OHT (+) but without APH (-) (RAS). (E-G) DNA combing analysis of ER:RAS FSE-hTert cells with (+) or without (-) 4-OHT and APH treatments as indicated. Individual replication fork rates (kb/min); at least 270 fibers per condition were analyzed (E). Individual fork distances (kb); at least 130 forks per condition were analyzed (F). Individual sister fork symmetry ratios; at least 110 forks per condition were analyzed (G). Red lines indicate medians. (G) Dashed green line indicates asymmetry ratio threshold. Data are the summary of two independent experiments. *P* values calculated by one-way ANOVA (E-G) and by Mann Whitney rank-sum test (D), ns - nonsignificant; * *P* < 0.05, ** *P* < 0.01, *** *P* < 0.001, **** *P* < 0.0001. Scale bars, 10 μm.

**Supplementary Figure 6. Related to Figure 4. RNA-seq analysis of RAS expressing cells.** (**A**) Principal component analysis (PCA) plot of gene expression data showing the 1st and 2nd principal components for control (red), RAS day 2 (green) and RAS day 4 (blue). (**B**) Hierarchical clustering heat map for differentially expressed genes (DEGs) between RAS expressing cells for 2 or 4 days (RAS day 2 and RAS day 4, respectively) and control cells. The Z-score centered log2-transformed gene in each sample is presented using a color scale. Three independent biological replicates are presented (1-3). (**C**) Top: Venn diagrams of upregulated (UP) or downregulated (DOWN) DEGs after RAS activation, showing overlap between RAS day 2 and RAS day 4. Bottom: Gene ontology (GO) term analyses of DEGs shared by RAS day 2 and RAS day 4. The top 10 enriched biological processes are shown.

**Supplementary Figure 7. Related to Figure 4. TOP1 downregulation generates classical replication stress.** (**A**) RNA-seq and RT-qPCR analyses of TOP1 in ER:RAS FSE-hTert cells treated with 4-OHT for 2 and 4 days (RAS day 2 and RAS day 4, respectively) relative to non-treated cells. For the RT-qPCR analysis, the values were normalized to those of RNA polymerase II and are the average fold change (mean ± s.e.m, n = 3). (**B**) Protein levels of TOP1 and GAPDH in ER:RAS FSE-hTert cells without 4-OHT treatment (Control), treated with two independent siRNAs against TOP1 (siTOP1-1 and siTOP1-2) and non-targeting siRNA (siCtrl), as indicated. (**C-E**) DNA combing analysis of ER:RAS FSE-hTert cells without 4-OHT (control), treated with two independent siRNA against TOP1 (siTOP1-1 and siTOP1-2) and non-targeting siRNA (siCtrl), as indicated. Individual replication fork rates (kb/min); at least 280 fibers per condition were analyzed (**C**). Individual fork distances (kb); at least 220 forks per condition were analyzed (**D**). Individual sister fork symmetry ratios; at least 120 forks per condition were analyzed (**E**). Red lines indicate medians. (**E**) Dashed green line indicates asymmetry ratio threshold. Data are the summary of two independent experiments. (**F**) Percent of cells with indicated number of co-localized γH2AX and 53BP1 foci in ER:RAS FSE-hTert cells without 4-OHT treatment (Control), treated with two independent siRNAs against TOP1 (siTOP1-1 and siTOP1-2) and non-targeting siRNA (siCtrl), as indicated. At least 400 nuclei per condition were analyzed. Data are the summary of two independent experiments. *P* values calculated by one-way ANOVA. ns – nonsignificant, ** *P* < 0.01, *** *P* < 0.001, **** *P* < 0.0001.

**Supplementary Figure 8. Elevated expression of TOP1 and RAS correlate in cancers.** (**A**) Expression levels (TPM) of TOP1 in tumor samples (red) and tissue matching normal samples (green) from 9 cancers, as indicated. (**B**) Scatter plots of expression (TPM) correlation between TOP1 and RAS normalized to GAPDH in tumor samples, as indicated. Pearson’s correlation coefficient is presented.

**Supplementary Figure 9. R-loop restoration underlies replication rescue.** (**A**) Protein levels of TOP1 and GAPDH in ER:RAS FSE-hTert cells with (RAS) or without (Control) 4-OHT treatment and HU and APH treatments as indicated. (**B**) Dot plot of mean nuclear fluorescence intensity of RNA-DNA hybrid specific antibody (S9.6) in individual nuclei in ER:RAS FSE hTert cells with (RAS) or without (Control) 4-OHT treatment, siRNA treatment or HU/APH treatments as indicated. Data for Control and RAS-cells with siRNA treatment are as presented in Figure 6H.

**Supplementary Figure 10. Model for TOP1-dependent accelerated replication and DNA damage.** Model of how RAS induces replicative stress generating DNA damage in pre-senescent cells. RAS activation elevates TOP1 expression, which reduces R-loop levels generating accelerated DNA replication rate progression resulting in DNA damage accumulation.

**Supplementary Table 1. Related to Figure 4. Expression fold change of replication annotated genes.** Expression level of replication annotated genes (GO:0006260) and TBP and ISG15, averaged normalized counts and log2 of the fold change are shown.

