## Supplementary Figures for "Topoisomerase 1 dependent R-loop deficiency drives accelerated replication and genomic instability"

### Supplementary Figure 1

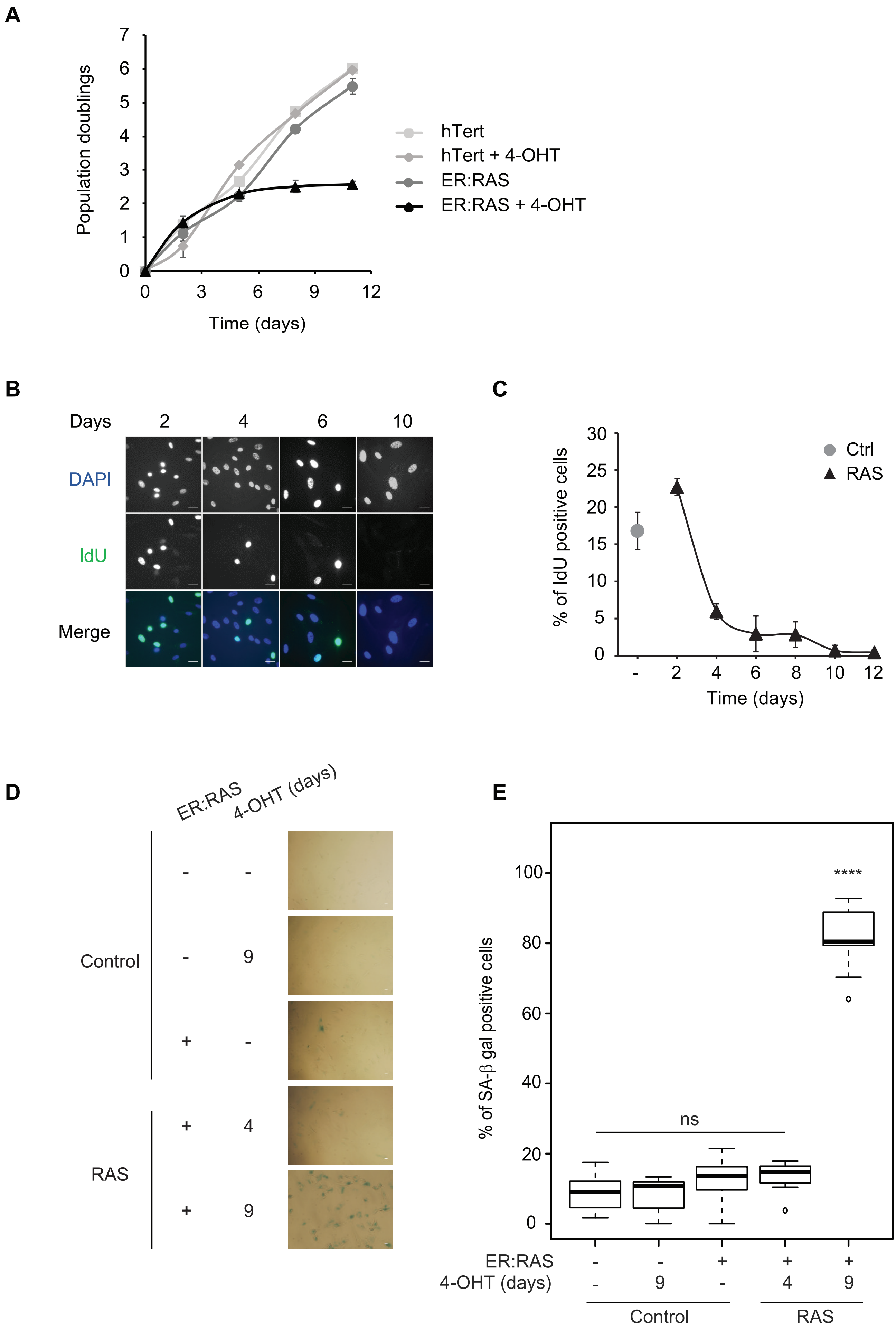

#### Supplementary Figure 2

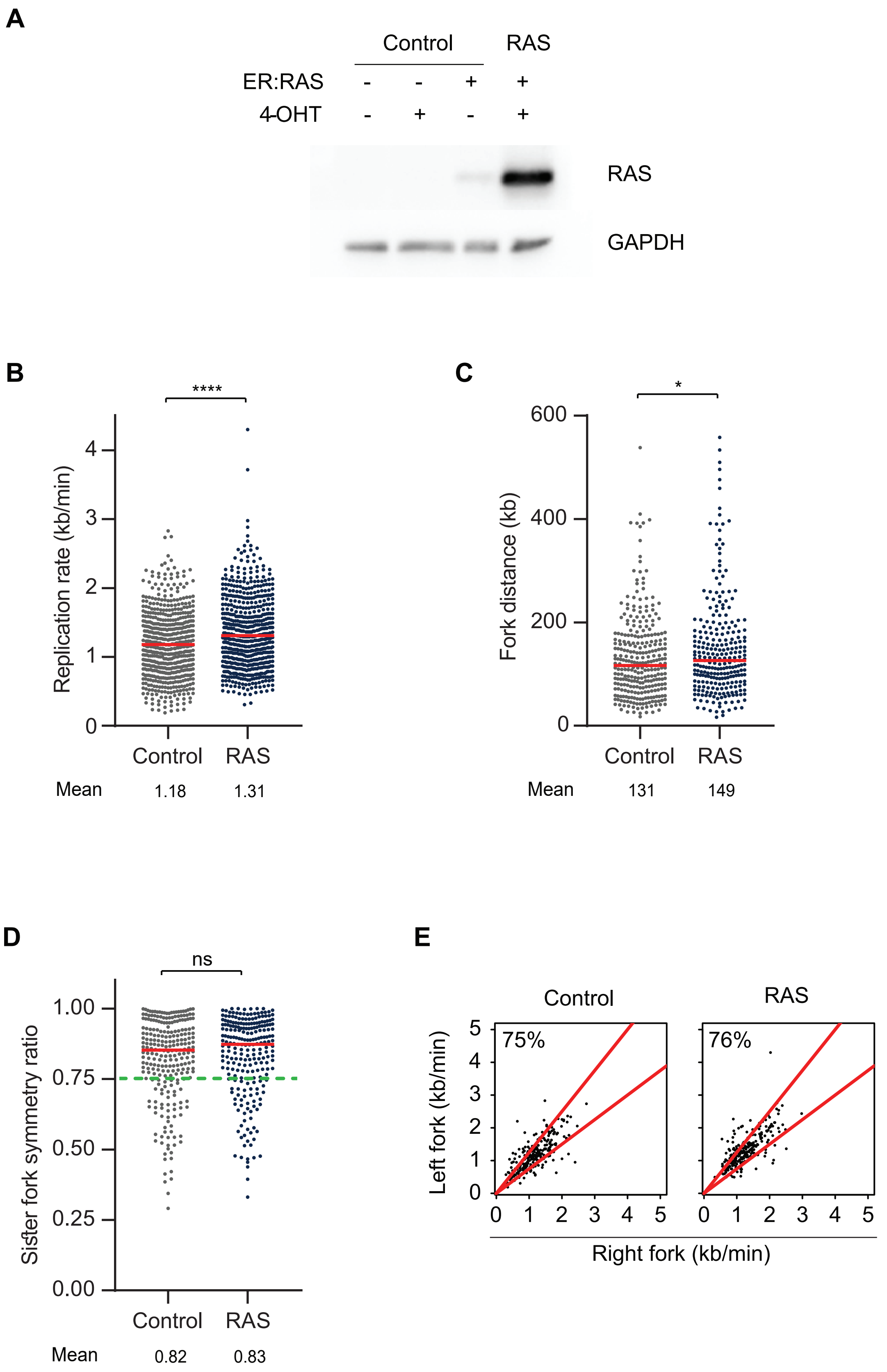

#### Supplementary Figure 3

A

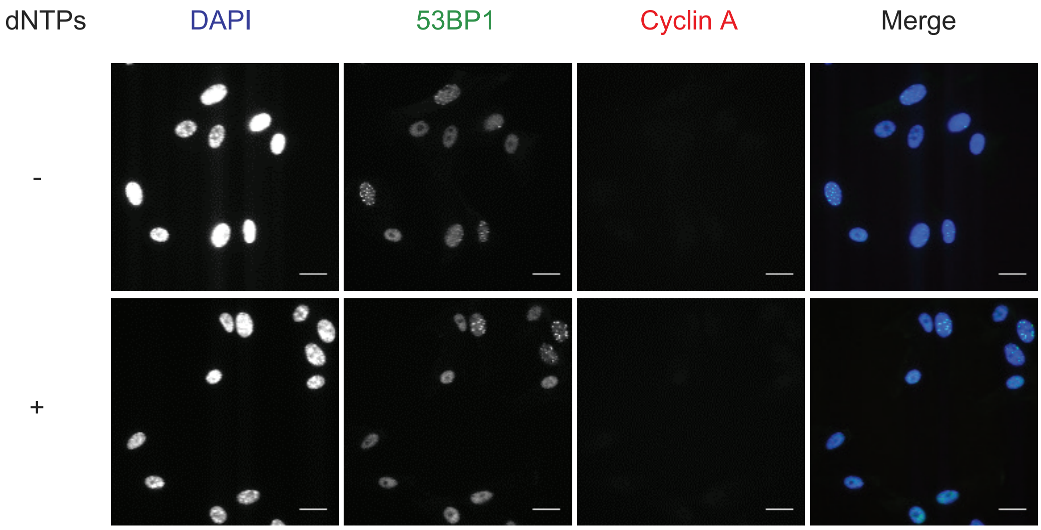


B

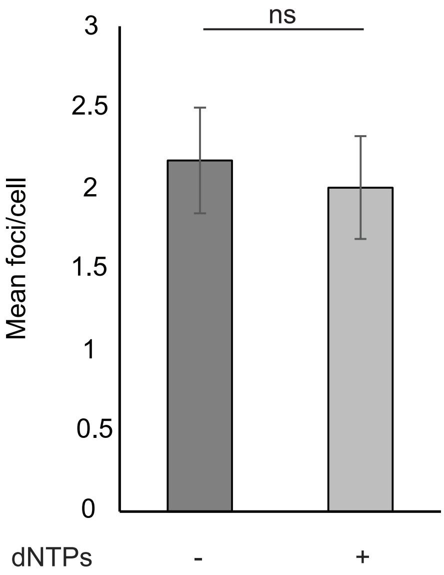

#### Supplementary Figure 4

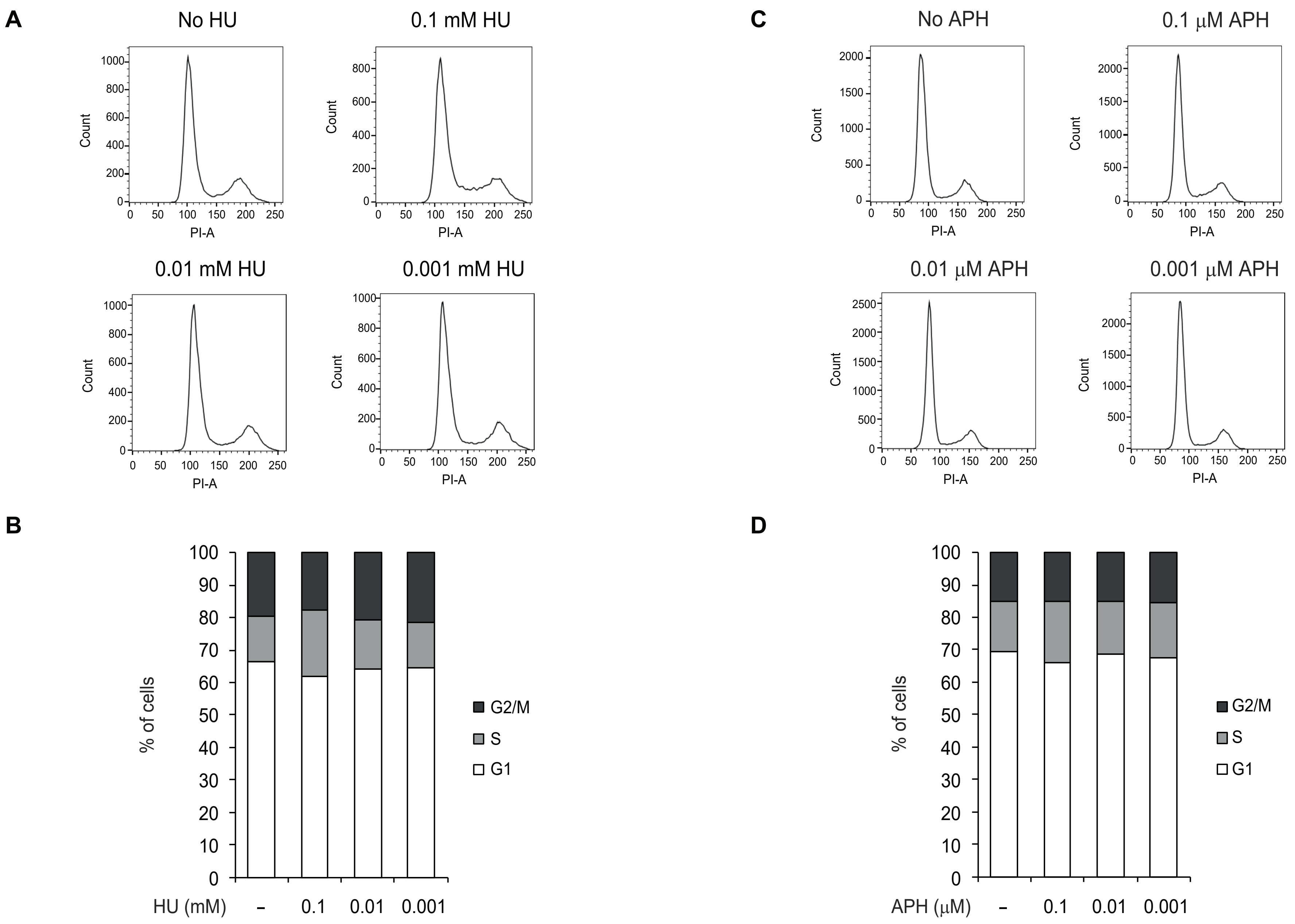

#### Supplementary Figure 5

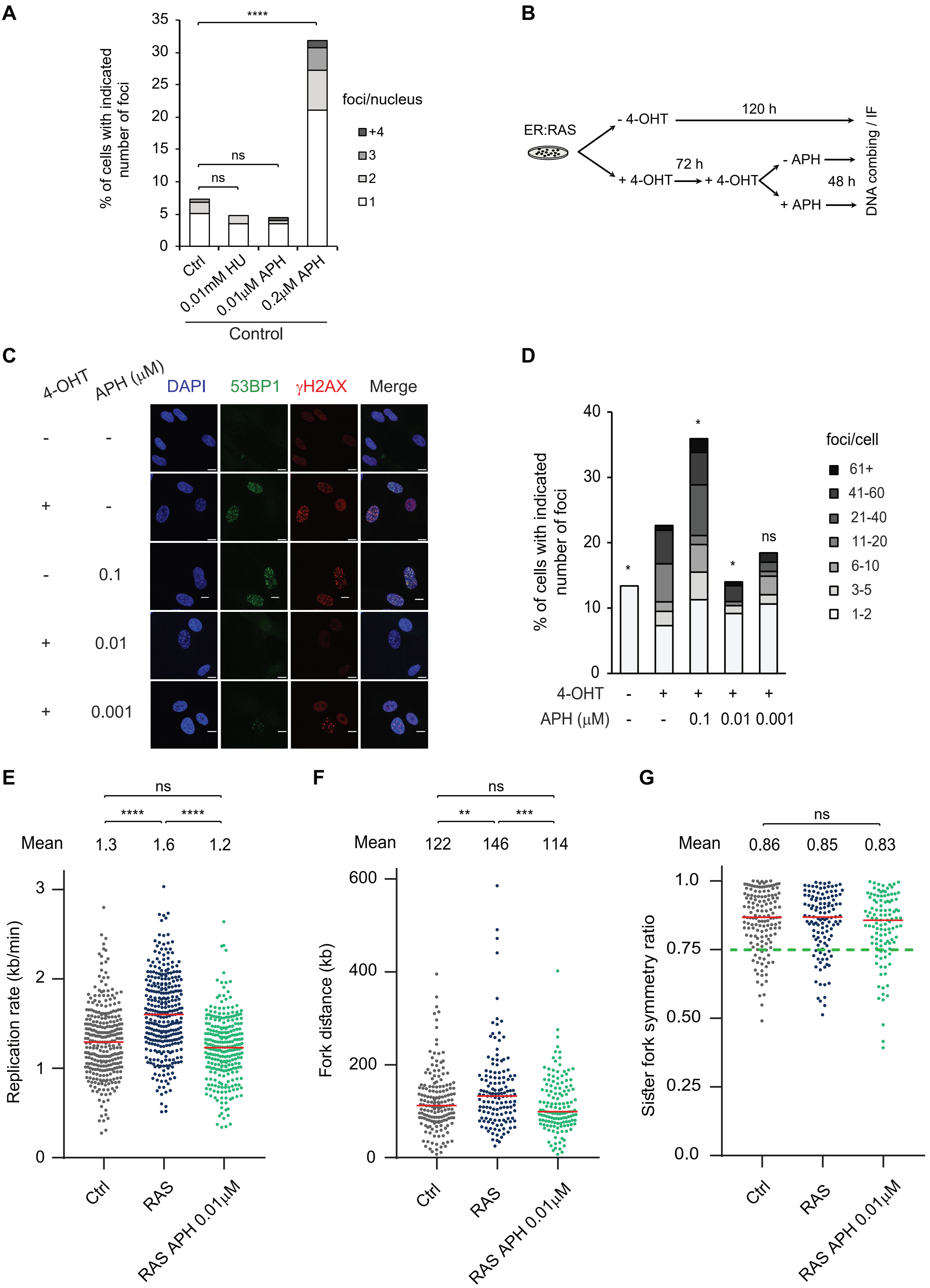

### Supplementary Figure 6

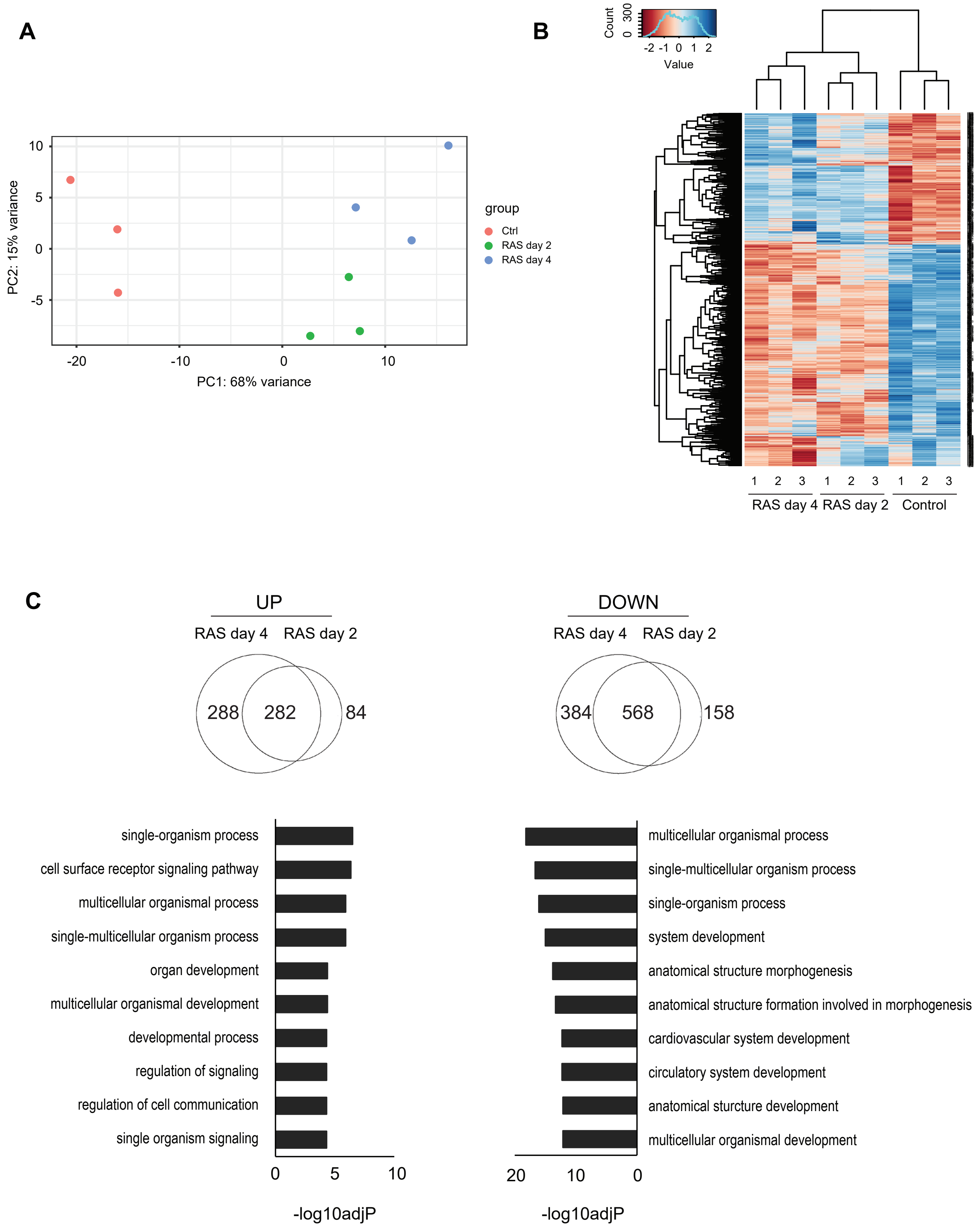

### Supplementary Figure 7

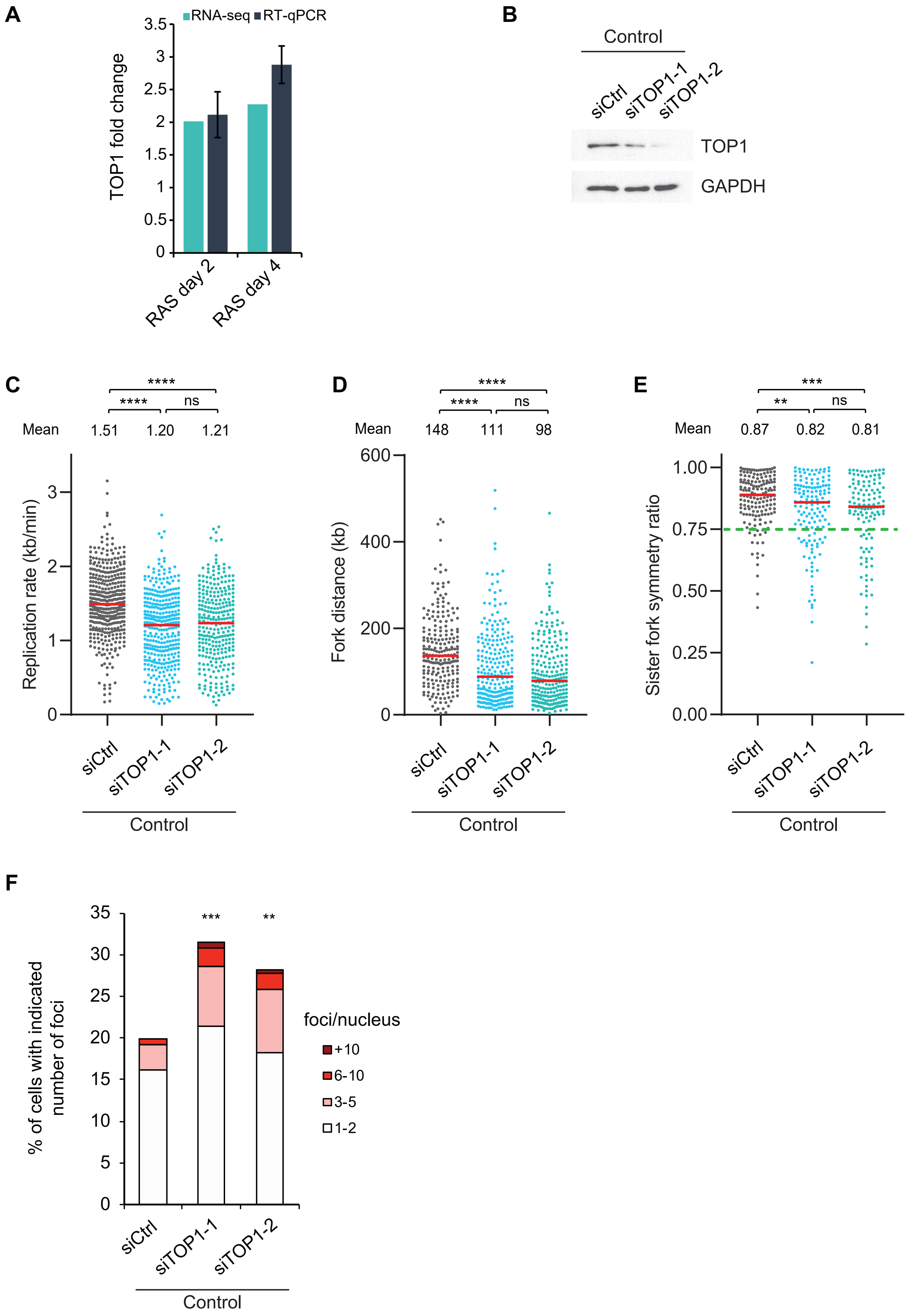

**Supplementary Figure 8**

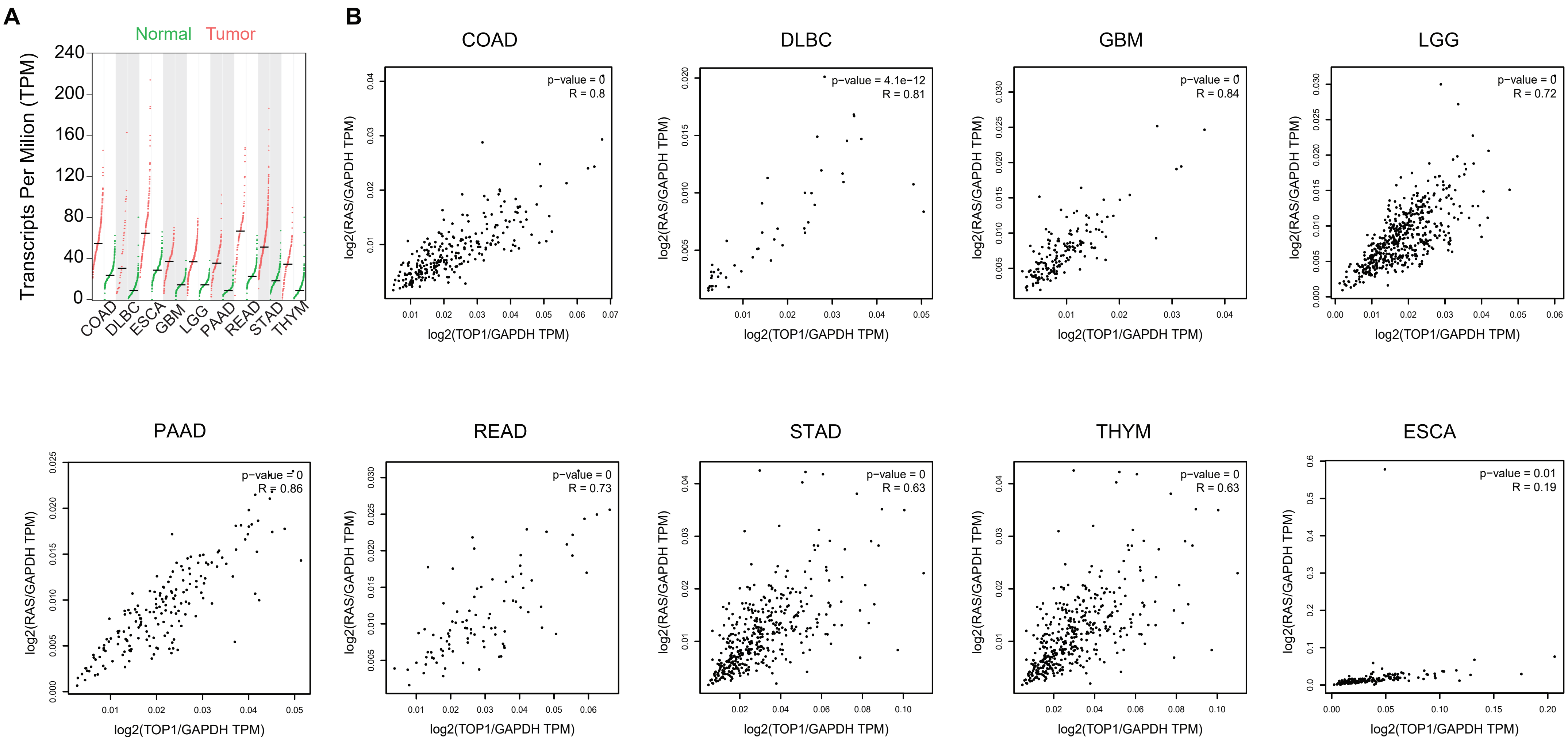

#### Supplementary Figure 9

**A**

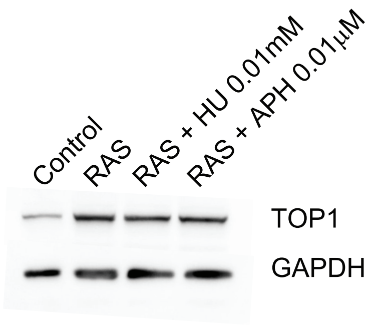


**B**

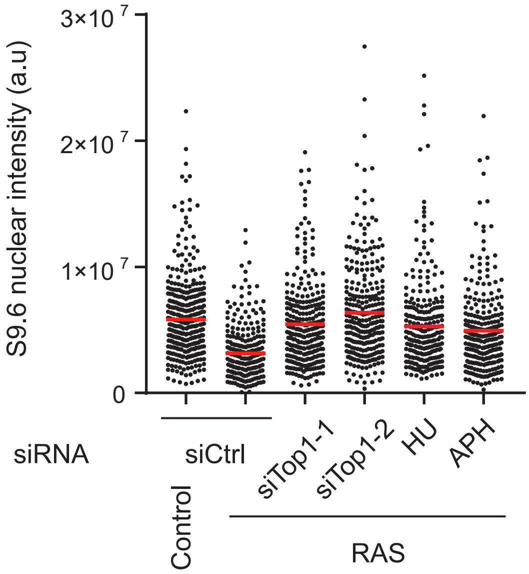

#### Supplementary Figure 10

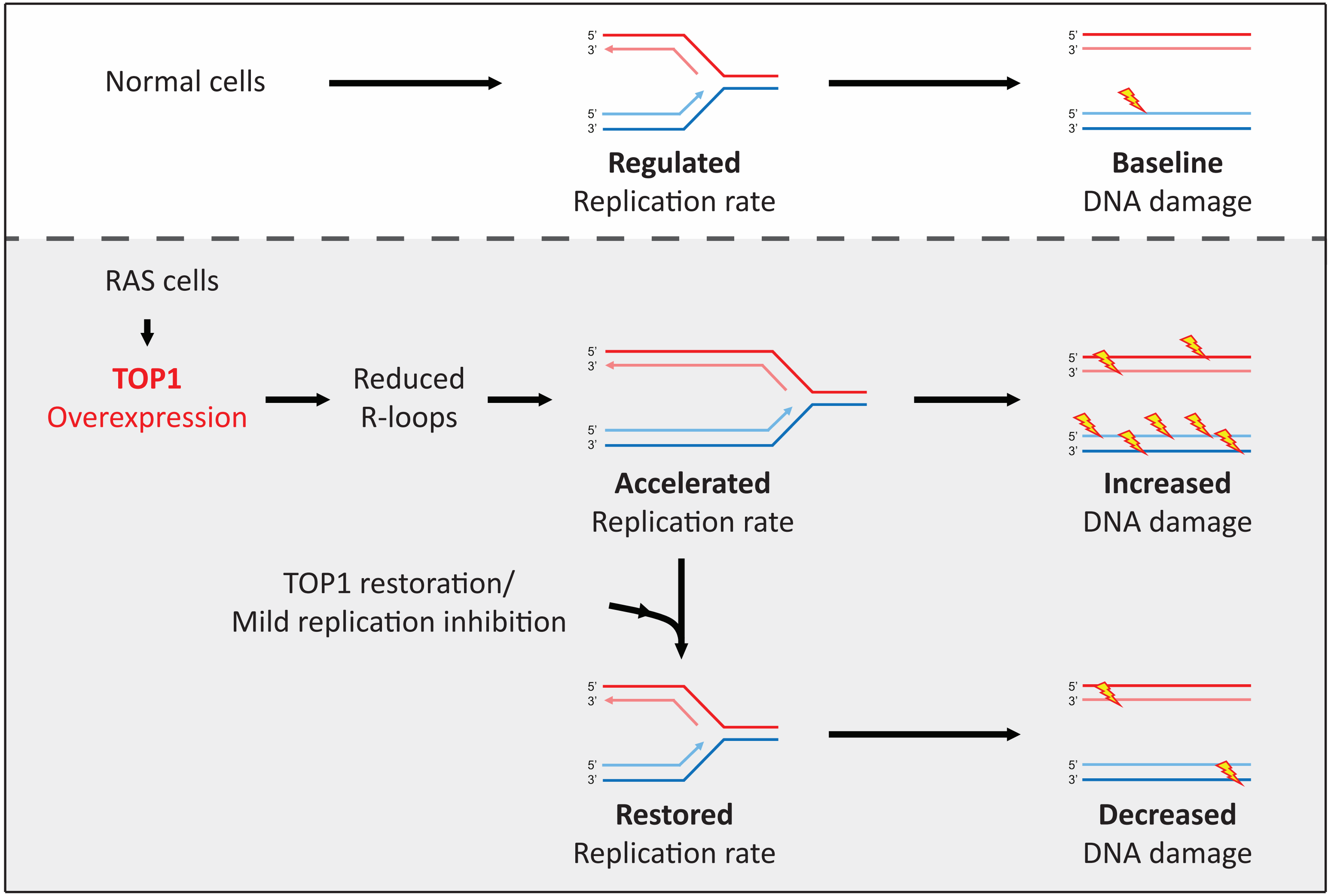
